# RABV L protein plays a role in immune escape through its methyltransferase activity

**DOI:** 10.1101/2025.09.12.675814

**Authors:** Marilys Castet, Wahiba Aouadi, Lauriane Kergoat, Ambre Bulteau, Rachel Legendre, Rania Ouazahrou, Lena Feige, Hugo Varet, Aurore Attina, Christophe Hirtz, Alexandre David, Etienne Decroly, Hervé Bourhy, Florence Larrous

**Author notes:** Co-first author (Marilys Castet, Wahiba Aouadi).

## Abstract

Viruses in the *Mononegavirales* order encode a large protein that orchestrates replication, transcription, and the capping of viral RNA. This protein, comprising over 2.000 amino acids, contains an RNA-dependent RNA polymerase, a capping domain, and a methyltransferase (MTase) domain involved in methylating the cap structure. The MTase domain features a conserved K-D-K-E catalytic tetrad -typical of 2′O-methyltransferases-which is essential for methylating viral mRNA caps at both the N7 and 2′O positions. However, the role of these residues in other epitranscriptomic modifications of rabies virus (RABV) RNAs remains poorly characterized.

To further explore the role of mRNA cap methylation in the immune evasion strategies of RABV, we investigated the functional contribution of the K-D-K-E motif within the MTase domain, using the Thai isolate as a model. Using reverse genetics, we demonstrated that the mutation K1830R in the K-D-K-E tetrad of the Tha MTase domain induces changes in the methylation landscape of viral mRNAs and, intriguingly, of host mRNAs. In addition, viruses harbouring the K1830R mutation are more sensitive to interferon-α and exhibit a less pathogenic phenotype *in vitro* and *in vivo* compared to the wild-type virus.

Overall, these results suggest that the regulation of viral and cellular RNA methylation landscapes plays a crucial role in controlling RABV infection. Although the exact role of these epitranscriptomic modifications is not yet fully understood, some of these methylations appear to have proviral effects and enhance viral propagation by allowing RABV to efficiently evade the host’s antiviral response.

**Importance:** This study highlights the pivotal role of the K-D-K-E catalytic domain included in the methyltransferase domain of the large protein of Rabies virus, by modelling viral RNAs with epitranscriptomic changes. For the first time, we identify specific methylations on the viral RNA, such as 2’-O and m6A methylations, which seem to enable the virus to mask its RNA and evade detection by the host’s pattern recognition receptors. These epitranscriptomic modifications affect not only viral RNAs but also cellular RNAs, underscoring a complex interplay between viral and host mechanisms. We further demonstrate that RABV harbouring an altered K-D-<u>R</u>-E catalytic domain, exhibit differential methylation patterns correlated with increased sensitivity to IFN and lower pathogenicity. This emphasizes the importance of this domain in virulence and immune evasion.

## Introduction

The order *Mononegavirales* (non-segmented negative-strand RNA viruses) contains important human viruses, including RABV, measles virus, respiratory syncytial virus (RSV) and Ebola virus. RABV is one of the most pathogenic *Mononegavirales*, causing about 59.000 human deaths per year (1). As RABV propagation is linked to an inefficient initial antiviral response, it is thus imperative to understand how this virus escapes early detection by the innate immune system and thus prevents innate immunity triggering. These mechanisms are also key in controlling neuronal infections, as the immune system is less effective in virus clearing beyond the blood–brain barrier.

Indeed, the innate immune system is the first barrier against viral infections, relying on the recognition of foreign nucleic acids and the rapid induction of interferon (IFN) pathways. In recent years, epitranscriptomic modifications of RNA, particularly methylation events, have emerged as key regulators of antiviral defense, shaping both host transcriptional responses and direct sensing of negative-strand RNA viruses(2, 3, 4, 5, 6). Methylations can target different nucleotide atoms (N7, 2’O, N6, N1, C5, etc.) (7–10) and either the cap structure decorating the 5’ ends of mRNA (11, 12) or nucleotides within the RNA chain (13, 14). Although it is well known that N7 methylation of the cap structure is essential for mRNA translation into viral proteins (11), the role of 2’O methylation of the cap structure was only recently evidenced as a cell self-marker (15–17).

Since *Mononegavirales* do not have access to the host mRNA capping machinery, which is located in the nucleus, they encode their own capping machinery. Indeed, their replication/transcription steps are ensured by the viral Large protein (L) (18, 19). The multifunctional enzyme L is a single polypeptide of more than 2,000 amino acid residues that carries all the enzymatic activities required for the transcription and replication of viral mRNA (20, 21). Thus, the L protein contains three main parts: i) the RNA-dependent RNA polymerase (RdRp), ii) the capping domain (CAP), and iii) the MTase domain (Figure 1A). The CAP domain is a GDP polyribonucleotidyl transferase (PRNTase) that transfers GDP to a newly formed 5’ monophosphate RNA (18, 19, 22). The MTase domain next methylates the 2’OH ribose of the first transcribed nucleotide and the N7 position of the guanylate cap structure (11).The 2’O MTase domain of RABV, Vesicular stomatitis virus (VSV), and all the other members of the order *Mononegavirales* share a common structural organisation with the highly conserved K-D-K-E tetrad of 2’O MTase (18, 21, 23–25) (Figure 1A-C). Site-directed mutagenesis performed on the RABV MTase provided functional importance of the K-D-K-E motifs (25). These mutants showed severely impaired viral replication and strong attenuation *in vivo*, with a notably increased sensitivity to type I IFN responses. These data confirm the dual role of the K-D-K-E motif in viral replication and immune evasion (25). However, the exact role of RABV catalytic K-D-K-E MTase residues in epitranscriptomic modifications of RABV and cellular RNAs remains poorly characterized.

**Figure 1.**
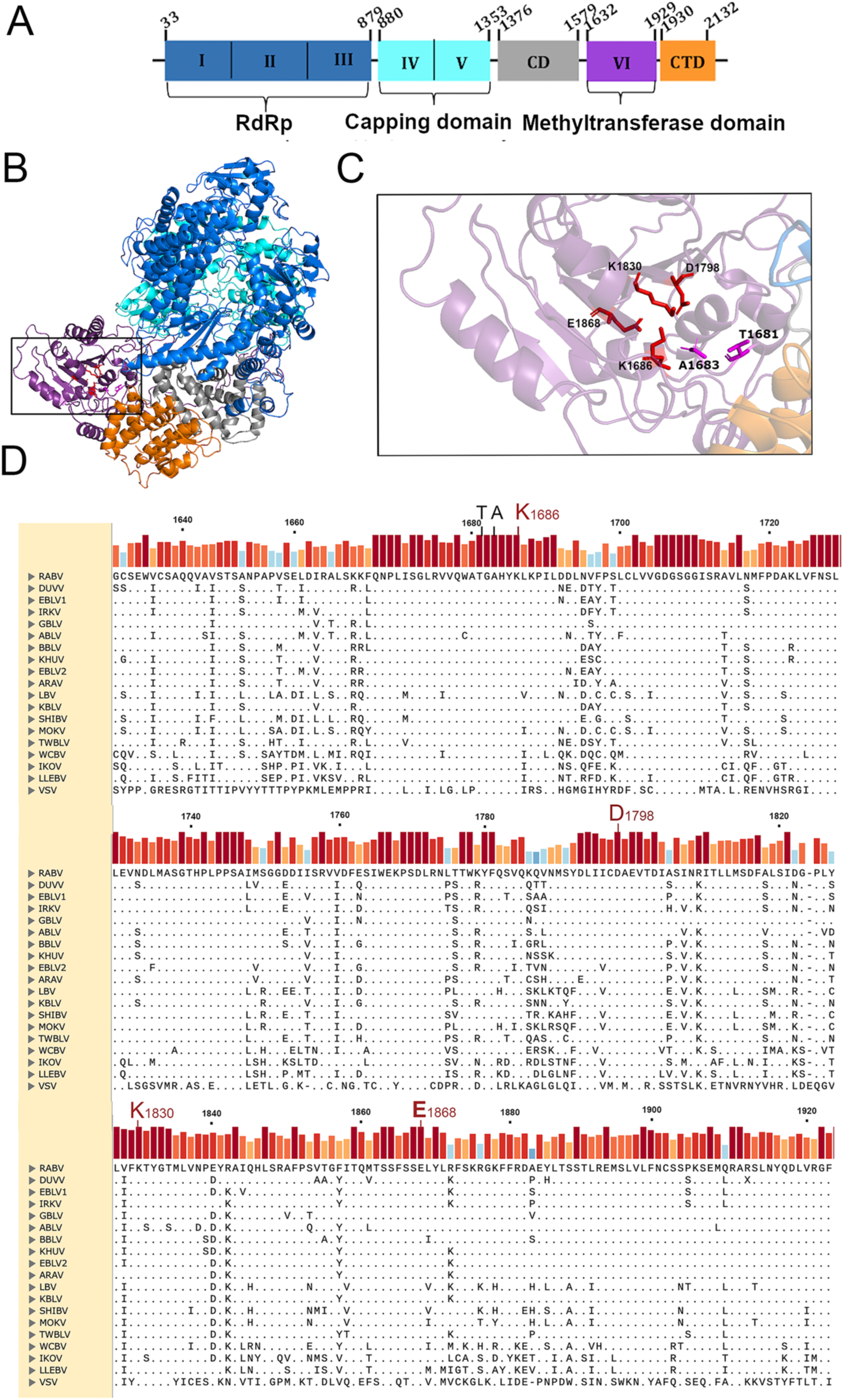
Characterisation of the methyltransferase of rabies virus. **(A)** Schematic representation of the L polymerase from rabies virus (RABV), showing its multi-domain architecture. The structure is color-coded by domain as follows: (i) **RNA-dependent RNA polymerase (RdRp) domain** (*blue*): catalyses viral RNA synthesis and adopts the typical “right-hand” configuration with fingers, palm, and thumb subdomains. (ii) **Capping (Cap) domain** (*turquoise*): facilitates the addition of the 5′ cap to nascent viral mRNAs, a crucial modification for RNA stability and translation. (iii) **Connector domain (CD)** (*grey*): serves as a structural bridge between the Cap and MT domains, likely playing a role in domain positioning and flexibility. **(iv) Methyltransferase domain (MT)** (*purple*): catalyses methylation of the mRNA cap structure. Its catalytic core includes the conserved **K-D-K-E** motif (Lys–Asp–Lys– Glu), essential for both guanine-N7 and 2′-O methylation activities. **(v) C-terminal domain (CTD)** (*orange*): may contribute to interactions with the phosphoprotein (P) and assist in coordinating transcription and replication. Conserved regions throughout NNS RNA virus L proteins are indicated by I, II, III, IV, V, and VI. **(B)** Structural model of the RABV L (ribbon structure). The model was built by alignment of RABV L and vesicular stomatitis virus (VSV) (PDB entry PDB:5A22) sequences using the Swiss model and the PyMOL software. **(C)** The zoom enlargement focuses on the catalytic core of the methyltransferase domain, highlighting the **K-D-K-E** motif, which is strictly conserved among lyssaviruses and critical for enzymatic methylation of viral mRNA caps. **(D)** Amino acid comparison of the **MT domain** from representative lyssaviruses and vesiculovirus, showing strong conservation of the K-D-K-E catalytic motif and associated active-site architecture. The alignment of amino acid sequences was performed with Clustal Omega in Snappgene version 8.0.3 on 19 different isolates. *Lyssavirus* Aravan (ARAV), EF614259 ; *Lyssavirus australis* (ABLV), <u>AF081020</u>; *Lyssavirus bokeloh* (BBLV), <u>JF311903</u> ; *Lyssavirus caucasicus* (WCBV), <u>EF614258</u> ; *Lyssavirus Duvenhage* (DUVV), <u>EU293119</u> ; *Lyssavirus formosa* (TWBLV), <u>MF472710</u> ; *Lyssavirus gannoruwa* (GBLV), <u>KU244266</u> ; *Lyssavirus hamburg* (EBLV1) MF187859. ; *Lyssavirus helsinki* (EBLV2), <u>EF157977</u>; *Lyssavirus ikoma* (IKOV), <u>JX193798</u> ; *Lyssavirus irkut* (IRKV), <u>EF614260</u>; *Lyssavirus khujand* (KHUV), <u>EF614261</u> ; *Lyssavirus kotalahti* (KBLV) <u>LR994545</u> ; *Lyssavirus lagos* (LBV), <u>EU293108</u>; *Lyssavirus lleida* (LLEBV), <u>KY006983</u>; *Lyssavirus mokola* (MOKV), EU293118 ; *Lyssavirus rabies (RABV),* EU293121 *; Lyssavirus shimoni (*SHIBV), GU170201; *Vesiculovirus newjersey* (VSV), <u>JX121109.</u>

To further explore the role of mRNA cap methylation in the immune evasion strategies of RABV, we investigated the functional contribution of the K-D-K-E catalytic motif in the MTase domain of the L protein, using a field isolate, Tha strain, as a model. This particular strain is known for its ability to suppress NF-κB and JAK-STAT signalling pathways (26) via its phosphoprotein (P) (27, 28) and matrix protein (M) (29, 30), providing a robust experimental model to dissect the contribution of L-mediated RNA methylation in modulating host responses.

We generated site-specific mutants within the K-D-K-E motif, focusing on the K1830R substitution (conserved lysine to arginine) in the MTase domain. This mutant virus remained viable, enabling detailed phenotypic analysis. *In vitro* studies demonstrated increased sensitivity to type I interferon (IFN-I) and attenuated replication, while *in vivo* studies revealed a significant loss of pathogenicity, indicating the critical role of K1830 in immune evasion and RABV virulence. Quantitative LC-MS/MS analysis of polyadenylated RNA from infected cells revealed differential methylation levels, including changes in 2′-O-methylation (Nm) and N6-methyladenosine (m⁶A). These findings were validated using sequencing-based methods: RiboMethSeq for mapping 2′-O-methylation sites and nanopore RNA sequencing for detecting m⁶A modifications in viral and host mRNAs.

Together, these approaches aim to explore the interplay between viral MTase activity, host RNA modification pathways, and the antiviral IFN response. Our study provides new insights into how mutations in the K-D-K-E motif of RABV MTase affect not only viral immune evasion and pathogenicity but also modulate the host epitranscriptome during RABV infection of neurons.

## Results

It is currently accepted that the MTase domain of the L protein, which is highly conserved within the order *Mononegavirales*, is responsible for the 2’O and N7 methylations of the viral messenger RNA (mRNA) cap structure. However, the role of RABV catalytic K-D-K-E residues in other epitranscriptomic modifications of RABV and cellular RNAs remains elusive. To decipher the roles of the viral MTase on viral and/or cellular RNAs, we introduced several mutations in the L protein gene of Tha virus: residues K1686, D1798, K1830 and E1868, which are involved in this K-D-K-E 2’O-MTase catalytic tetrad, as well as T1681, a residue located at the beginning of the MTase domain (located outside of the catalytic tetrad), which was used as a control (Figure 1A-D).

### A mutation in the K-D-K-E motif of the RABV MT domain limits viral escape to type I interferons

To investigate whether mutations in the K-D-K-E motif of the L protein could account for a defect in viral replication, we introduced single mutations either in L-expression plasmids (for minireplicon assays) or in the complete genome of the Tha strain by substituting residues of the tetrad K1686, D1798, K1830, E1868 and the control T1681, by R, N, R, N and S residues respectively. K was substituted with R residues to maintain a basic amino acid. D and E are diacid amino acids and they were both substituted with N, an amidated amino acid. T residue was substituted by another sulphur-containing amino acid S. The viruses containing L protein mutations were named Tha-L_K1686R_, Tha-L_D1798N,_ and Tha-L_K1830R_, Tha-L _E1868N_, and Tha-L_T1681S_ according to their amino acid substitutions (Figure 2A).

**Figure 2.**
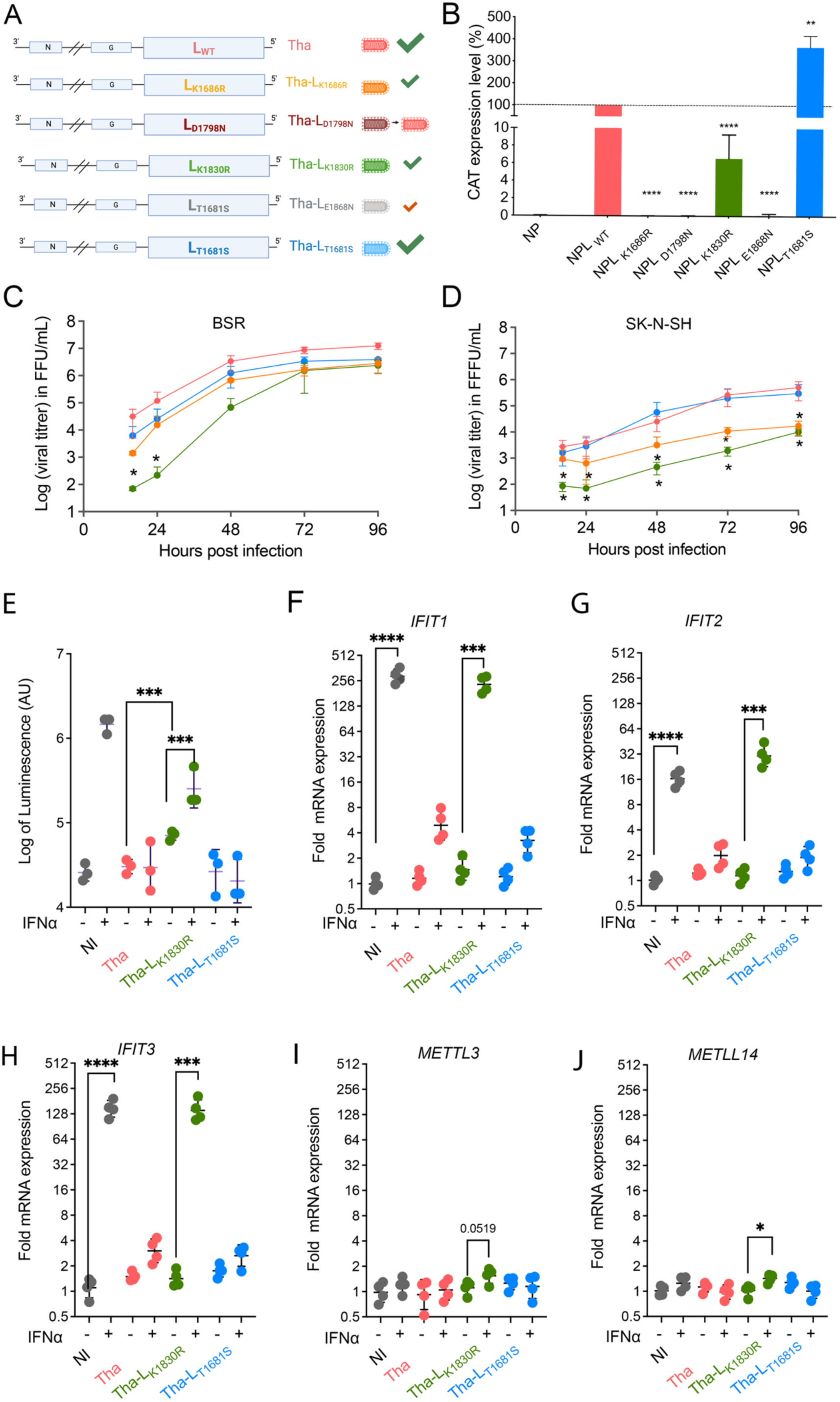
Mutations on the K-D-K-E motif of RABV MT domain affect rabies infection and modify cell response. **(A)** Introduction of a single mutation from the K-D-K-E motif of the viral MTase in the complete genome of the Tha virus. The mutated residues are indicated on the schematic representation of the genome. In pink, recombinant wild-type Tha virus; in orange, recombinant Tha virus mutated on residue K in position 1686 of the L protein mutated in R (named Tha-L_K1686R_); in burgundy, Tha-L_D1798N_; in green, Tha-L_K1830R_; in grey, Tha-L_E1869N_; in blue, Tha-L_T1681S._ A green tick indicates an efficient rescue. **(B)** Minireplicon assay with mutated L protein. The activity is normalized according to the CAT activity obtained with the wild-type L protein. The results represent the mean ± SEM of at least three independent experiments. Significant results are indicated by stars (p<0.05). **(C-D)** Viral growth of the recombinant mutated viruses (Tha, Tha-L_K1686R_, Tha-L_K1830R,_ and Tha-L_T1681S_) in BSR and SK-N-SH cells. Viral titer was calculated at 16 hours post-infection (hpi), 24 hpi, 48 hpi, and 72 hpi. The results represent the mean ± SEM of at least two independent experiments. **(E)** Modulation of ISRE promoter activity in STING37 cells infected with Tha, Tha-L_K1686R_, Tha-L_K1830R,_ and Tha-L_T1681S_ viruses). The IFN pathway was exogenously activated 24 hours after infection using 1000 UI IFN-α for 24 h more (noted +) or untreated (noted -). Log of the absolute luminescence is registered and presented on the y-axis. The results represent the mean ± SEM of at least three independent experiments. Significant results are calculated using unpaired t-test with Welch’s correction (*: p<0.05; **: p<0.01; ***: p<0.001; ****: p<0.0001). **(F-J)** Quantification of gene expression of *IFIT1*, *IFIT2* and *IFIT 3*, *METTL3* and *METTL14* respectively. The IFN pathway was exogenously activated 24 hours after infection using 1000 UI IFN-α for 24 h more (noted +) or untreated (noted -). The fold mRNA expression is calculated relatively to non-treated, non-infected cells. Significant results are calculated using unpaired t-test with Welch’s correction (*: p<0.05; **: p<0.01; ***: p<0.001; ****: p<0.0001).

To study the impact of mutations introduced in the L protein on viral replication, we first performed a minireplicon assay (31), using the wild-type L plasmid or the mutated versions (Figure 2B). As expected, mutations in the residues of the catalytic tetrad drastically impaired replication activity compared to the wild-type L, whereas L_T1681S_ showed enhanced replication. Mutations L_K1686R_, L_D1798N_, and L_E1868N_ completely aborted the replication complex activity, whereas L_K1830R_ still induced a low level of activity (approximately 6% of the wild-type L activity). Then we investigated the capacity of recombinant viruses harbouring these mutations to be rescued. In agreement with the conserved replication activity of L_T1681S_, the Tha-L_T1681S_ could be rescued successfully (Figure 2A). Although a virus was rescued for Tha-L_D1798N_ (Figure 2A), the genome reverted immediately after the first passage to the wild-type sequence, indicating that the rescue was not successful. The mutation L_E1868N_ did not prevent production of virus Tha-L_E1868N,_ but its growth was heavily reduced compared to Tha virus (Figure 2A; Figure S2). Tha-L_E1868N_ could not be used in the rest of this work due to its low infectious titer.

While the K1686R and K1830R mutations initially appeared to strongly impact the activity of the replication complex, it was still possible to rescue viruses carrying those mutations that remained stable over time. However, the K1830R mutation alters the spread of the virus as seen by confocal analysis in Figure S3 compared to the wild-type Tha virus. The viral growth of the recombinant viruses, Tha, Tha-L_K1686R,_ Tha-L_K1830R,_ and Tha-L_T1681S_ was first studied on IFN-deficient BSR cells and then on the IFN-competent SK-N-SH cells (Figure 2C and D). As expected, Tha-L_T1681S_ shows a growth curve similar to that of the wild virus in both cell types. A mutation at residue K1686R impaired viral growth, as evident in our data for SK-N-SH cells, but had a minimal effect on viral growth in BSR cells 72 hours post-infection. However, Tha-L_K1830R_ was more significantly affected during the early phase of its growth on BSR cells (Figure 2C), and its replication was highly altered on SK-N-SH cells compared to BSR cells (Figure 2D). Thus, we next focused our study on the Tha-L_K1830R_, which appears to be highly affected in its growth, and on Tha-L_T1681S_, used here as a control that bears a mutation in the MTase domain without any modification of the viral growth phenotype.

### Mutations in the K-D-K-E motif of RABV MTase domain alter cell response

Since viral replication of Tha-L_K1830R_ is more impaired in IFN-competent cells such as SK-N-SH compared to immune-deficient BSR cells, we hypothesized that this mutation induced greater sensitivity of the virus to the host IFN pathway. More precisely, it has already been described that mutants lacking 2’O MTase activity induce higher expression of type I IFN and are more sensitive to type I IFN response (16) and that IFIT proteins can limit the RNA translation of mRNA lacking 2’O methylation (32).

To test the activation of the IFN pathway, STING37 cells expressing a luciferase reporter gene under ISRE promoter control were infected with Tha-L_K1830R_ and Tha-L_T1681S_ recombinant viruses. After 24 hours post-infection (hpi), IFN-α was added exogenously, the cells were lysed at 48 hpi, and the reporter activity was measured (Figure 2E). As expected, the luminescence activity measured for non-infected cells before IFN-α treatment was low, and IFN-α treatment induced expression of luciferase. Cells infected with Tha and Tha-L_T1681S_ revealed a similar fluorescent signals compared to the signal observed in uninfected cells, both before and after IFN-α treatment, in accordance with Tha’s ability to suppress NF-κB and JAK-STAT signalling pathways (26, 27, 28, 29, 30). In addition, cells infected with Tha-L_K1830R_ revealed a significantly higher luminescence signal than cells infected with the wild-type virus Tha. After IFN-α treatment, Tha-L_K1830R_-infected cells show an increase in luminescence (Figure 2E), as observed with the uninfected stimulated cells.

In parallel, we infected SK-N-SH cells with the same viruses, with or without exogenous IFN-α stimulation, and extracted cellular RNAs 48 hpi. Activation of the ISRE promoter was evaluated by measuring the expression of various Interferon-stimulated genes (ISGs) and particularly *IFIT1*, *IFIT2,* and *IFIT3* using the Taqman assay (Figure 2F-H). As expected, in the absence of exogenous IFN-α, Tha and Tha-L_T1681S_ escaped the host response and did not modulate expression of any of the afore mentioned *IFIT* genes (16). Surprisingly, Tha-L_K1830R_ also did not induce transcription of the ISG *IFIT1, 2,* and *3* in infected cells (Figure 2F-H). However, when IFN-α was added exogenously 24 hpi, a significant increase in the expression of these genes was observed only for Tha-L_K1830R_, as in uninfected cells. This result suggests that the virus mutated on residue K1830 of the catalytic tetrad of the MTase domain of the L protein cannot counteract the IFN type I response and is therefore more sensitive to IFN-α than the wild-type strain.

To further characterize the phenotype associated with Tha-L_K1830R_ infection, we analysed the relative gene expression of *METTL3* and *METTL14*, two genes coding for cellular methyltransferases involved in m6A labelling of cellular and/or viral mRNA. Gene expression of *METTL3* and *METTL14* on Tha-L_K1830R_ -infected cells showed a weak (1.5-fold) but significant induction upon IFN-α stimulation, conversely to other infections (Figure 2I-J), raising the possibility that the K1830R mutation of the MTase may influence host RNA modification pathways and induce feedback on the epitranscriptomic landscape during infection.

### Mutations in the K-D-K-E motif of the RABV MTase domain affect viral and cellular mRNA methylation levels

Our results indicate that mutations in the viral MTase catalytic domain (K1830R) regulate the expression of cellular MTases involved in the N6 methylation pathway. We therefore analysed the cellular mRNA methylation pattern after Tha, Tha-L_K1830R_ or Tha-L_T1681S_ infection. The cellular mRNAs were purified using Dynabeads before RNA hydrolysis by P1 nuclease, which hydrolyses RNA into nucleoside monophosphates in order to identify the nucleotides modification within the mRNA. We also uncapped the RNA using 5’ Pyrophosphohydrolase (RppH), in order to identify methylations on the cap structure (on guanosine N7 methylation) or on the first nucleotide of mRNA (methylations at 2’O, or N6 positions) (Figure 3). These digestions were analysed by mass spectrometry after HPLC separation (LC/MS-MS) (12, 14, 33) in order to identify the type of methylations and their positions on the nucleotide sequence (N6, 2’O or others) (Figure 4A) through HPLC separation.

**Figure 3.**
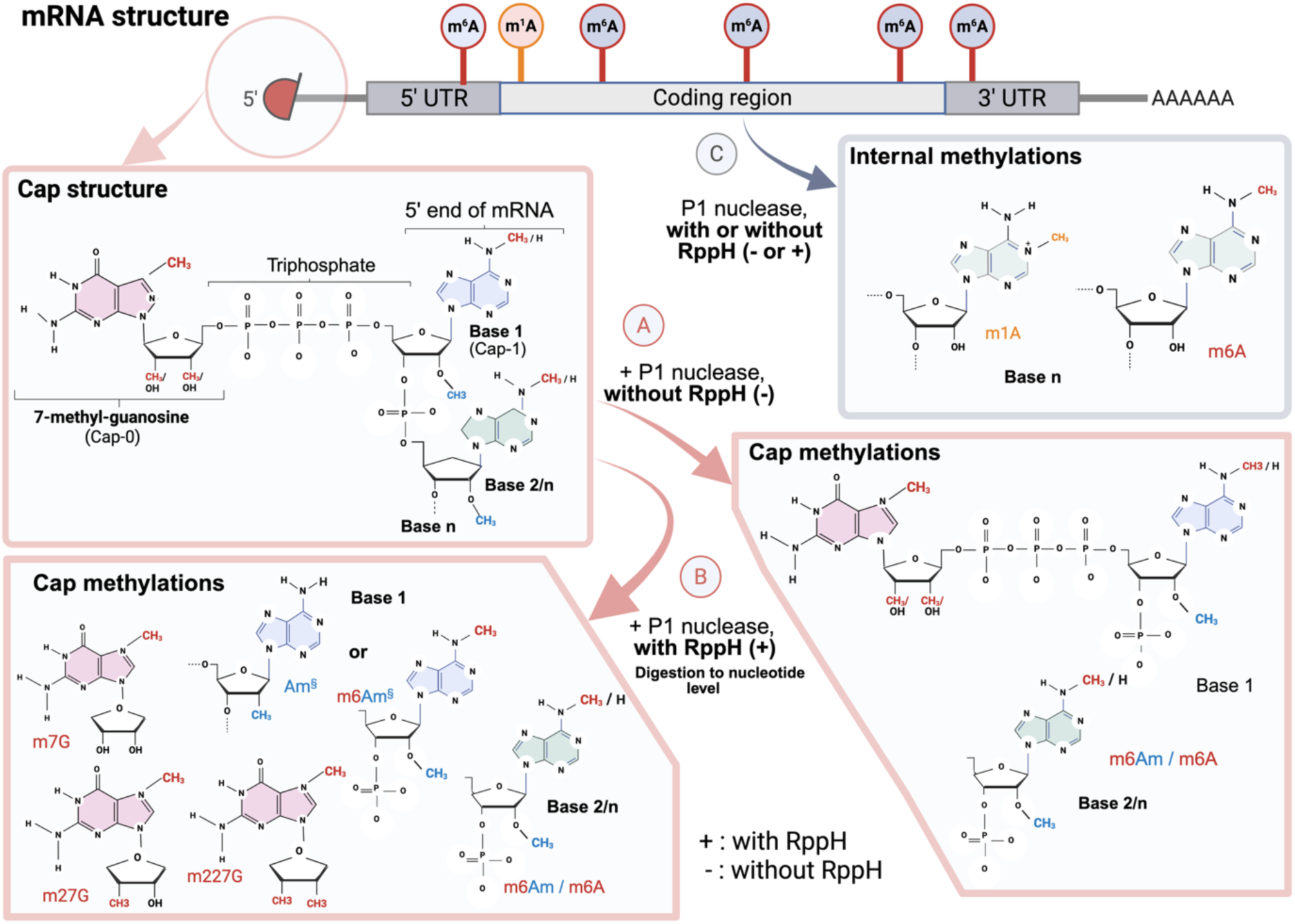
Schematic representation of the most representative modifications observed in our study on internal position of mRNA and in the cap structure of viral mRNA products. RppH is a decapping enzyme that cleaves the bond between the methyl-guanosine and the first base of mRNA. P1 nuclease cleaves only between each nucleoside of the RNA. **(A)** Treatment by P1 nuclease without RppH digestion will release nucleosides from the second base, which can be 2’O methylated on adenosine (Am) or on other nucleoside (substituted by Gm; Cm; Um according to the mRNA sequence, noted Am^§^ on the graph) or N6-2’O-methyl-adenosine (m6Am). **(B)** Treatment by P1 nuclease with RppH will totally digest the cap structure and release the m7G of the cap and the 2’O methylation on the first base (Am or Am^§^). N6-2’O-methyl-adenosine (m6Am) and N6-methyl-adenosine (m6A) of internal nucleoside will also be detected, but also single or double methylated at the N2 atom of the terminal guanosine (m2,7G and m2,2,7G, respectively). **(C)** P1 nuclease treatment with or without RppH digestion will release N6-methyl-adenosine (m6A) and N1-methyl-adenosine (m1A), which are in internal position of the mRNA.

**Figure 4.**
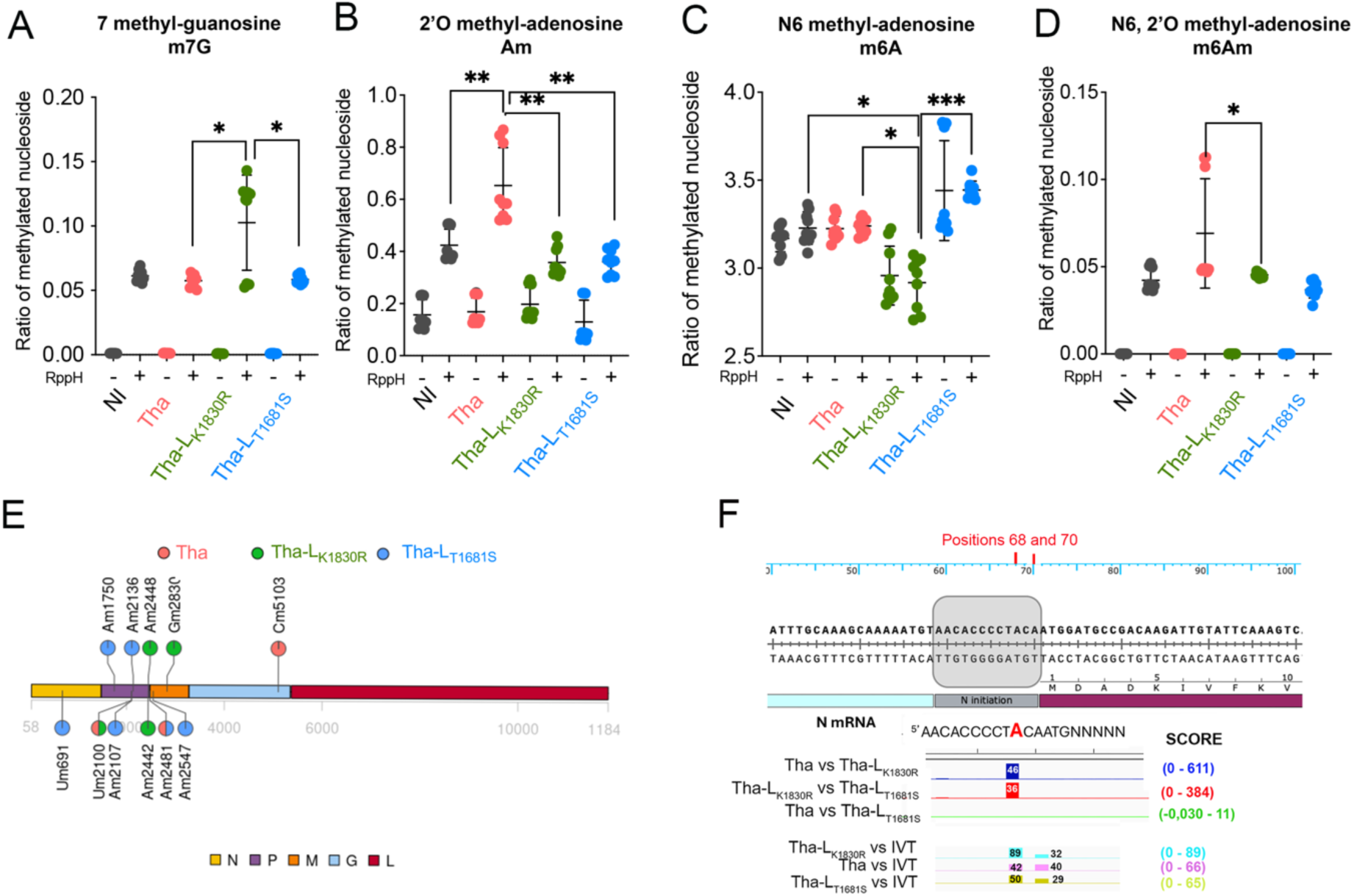
Mutations on the K-D-K-E motif of the RABV MT domain affect viral and cellular mRNA methylation levels. Quantitative LC-MS/MS measurements of 5′ terminal and internal modifications in poly(A) RNA fractions were performed on SK-N-SH cells infected with Tha, Tha-L_K1686R_, Tha-L_K1830R,_ and Tha-L_T1681S_. (**A-D)** Relative levels of 7 methyl-guanosine (m7G); 2’O-methyl-adenosine (Am); N6-methyl-adenosine (m6A) and N6-2’O-methyl-adenosine (m6mA) respectively, modifications obtained by digestion with (+) or without (-) RppH and P1 nuclease on purified mRNA from infected SK-N-SH cells (*n* = 3). The levels of nucleoside modifications were normalized to the total molar amount of canonical nucleosides N (N = C + U + G + A). Results are average over the levels of repbio. P-value adjustment was calculated by the Tukey method for comparing a family of 4 estimates. Significant results are indicated by star: *: p<0.05; **: p<0.01; ***: p<0,001; ****: p<0,0001. **(E)** Detection of 2’O methylation in internal position of viral mRNA from infected SK-N-SH. Modified residues identified by RibomethSeq are marked on the genome: in pink for Tha, in green for Tha-L_K1830R,_ and blue for Tha-L_T1681S_. Positions are considered as ribomethylated when they passed the score filters, conversely to the negative control (mixIVT). **(F)** Detection of m6A methylations in the internal position of viral mRNA from infected SK-N-SH. Position 68 on the Tha genome shows a differential methylation score between Tha-L_K1830R_ and Tha (in dark blue), between Tha-L_K1830R_ and Tha-L_T1681S_ (in red), and between Tha and Tha-L_T1681S_ (in green). Comparison between each virus and a mix of *in vitro transcript* (IVT): Tha vs IVT is indicated in pink, Tha-L_K1830R_ vs IVT in cyan, and Tha-L_T1681S_ vs IVT in light green. Positions 68 and 70 (in red) are on the 5’UTR of N mRNA (in grey). Differential methylation rates and scores between each comparison on bars. The heights of the bars on the graph are representative of the methylation scores for each residue.

Comparative analysis of the methylations rate upon P1 treatment only (without Rpph) and P1 treatment with additional Rpph treatment allowed us to segregate methylations that are only present on the cap (Guanosine or first nucleotide), that are only present in internal positions, or that are present on the cap and in internal positions. Methylations are only present on the cap when the rate is null without Rpph treatment (no methylation observed when uncapping), and positive with Rpph treatment. Methylations are only found in internal positions when the rate is similar with and without Rpph treatment (uncapping do not increase the methylation rate). Finally, methylations are present on the cap and in internal positions when the rate is positive without Rpph treatment (indicating internal methylations) and even higher with Rpph treatment (resulting in additional methylation due to uncapping), (Figure 3).

First, regarding the N7 methylation of the cap structure, the ratio of 7-methyl-guanosine (m^7^G) resulting from Cap-0 (methylation m7) and Cap-1 (2’O methylation of the first base) hydrolysis by RppH was similar in uninfected cells and cells infected with Tha and Tha-L_T1681S_ (Figure 3, 4A). Conversely, a significant increase in the m^7^G ratio is observed in cells infected with Tha- L_K1830R_ compared to the other types of infection (p<0.05). This significant increase is also observed for 2’O-7-methyl-guanosine (m^2,7^G) or 2’O-2’O-7-methyl-guanosine (m^2,2,7^G), which are respectively associated with single or double methylation at atom N2 of the guanosine. These methylations are present in smaller amounts on the Cap structure (Figure 4A; S4A-B). Additionally, we analyzed N6,2’-O-methyladenosine (m^6^Am) on the cap structure, showing that Tha L_K1830R_ infection causes a significant decrease in m^6^Am cap methylation levels compared to the wild-type virus (*p < 0.05, Figure 4D).

Regarding internal 2’O-methylation, similar low ratios were observed in infected or uninfected cells without RppH treatment (Figure 4B), suggesting that infection with Tha wild-type or mutants may not affect internal 2’O-methylation levels. However, the 2’O methylation (A_m_) at the first nucleotide of RNA (N1) (Cap-1), which shows an increased ratio after RppH digestion, leads to higher levels of A_m_ (Figure 3, 4B) following Tha infection. In contrast, Tha-L_K1830R_, Tha- L_T1681S_, or uninfected cells show similar 2’-O-methylation levels, thus suggesting that K1830 and T1681 residues may not be involved in methylation of the cap-1 structure.

We also analysed the internal mRNA methylations level by analysing methylation obtained in absence or in presence of RppH treatment (Figure 4B-C) and comparing the methylation with or without RppH treatment (Figure 3).

Regarding the internal N6 methylation of adenosine (m^6^A), levels were studied. Cellular infection with Tha or Tha-L_T1681S_ did not affect the m^6^A internal methylation levels, which are similar between Tha-infected and non-infected cells (Figure 4C). Interestingly, infection with Tha-L_K1830R_ caused a significant decrease in internal m^6^A methylation levels compared to the wild-type virus (*, p<0.05) or non-infected cellular RNAs (*, p<0.05), while Tha-L_T1681S_ caused a significant increase (***, p<0.001) in this type of methylation compared to Tha-L_K1830R_ (Figure 4C).

In addition, we also observed that Tha and Tha-L_T1681S_ induced a lower level of m^1^A (N1-methylation on adenosine) after RppH decapping, in comparison to uninfected cells or cells infected with Tha-L_K1830R_ (Figure S4E).

Other methylations were identified at lower frequency. Firstly, on the cap structure of the mRNA purified from infected cells, there was a significant decrease in 2’O methylation on C (C_m_) in

Tha-L_K1830R_-infected cell RNAs compared to Tha- (***: p<0.001) or non-infected cells (*: p<0.05, Figure S4C). Secondly, internal methylation levels were analysed on guanosine at the 2’OH position on G (G_m_), which revealed the same methylation ratios upon Tha-L_K1830R_ and Tha- L_T1681S_ compared to uninfected cells. Additionally, hypermethylation was observed in Tha-infected cell RNAs compared to non-infected ones (**: p<0.01, Figure S4D).

RABV infection, therefore, modulates the levels of methylation of mRNAs from infected cells, and the MTase domain of the L protein was directly or indirectly involved in these regulations, as infection by mutant virus Tha-L_K1830R_ induced more m^7^G, m^2,7^G, and m^2,2,7^G methylations on the Guanosine of the cap and less internal m^6^A methylations than infection by wildtype Tha virus.

Further analysis performed using RiboMethSeq or Nanopore, which will be described later, revealed that reads that aligned to viral RNAs represented only 2% to 20% of the total reads (Figure S6, Table S2). This, along with the high levels of methylation and the predominance of cellular RNAs in infected cells, strongly suggests that cellular mRNAs may also be influenced by virus-induced methylations.

### Modulation of 2’O methylation in internal positions of viral mRNA

To determine whether internal methylations were present on viral mRNA, we first analyzed the 2’O-methylation pattern of Tha, Tha-L_K1830R_ and Tha-L_T1681S_ mRNAs using RiboMethSeq technology (10). Purified mRNAs from infected cells were sequenced on the Illumina HiSeq 2500 platform to generate single-end 65 bp reads. We specifically focused on the reads that aligned to the rabies virus Tha reference genome (Lyssavirus rabies isolate 8743THA, accession number EU293121.1). The five viral mRNA (N, P, M, G, and L) were detectable (Figure S5A) with a coverage gradient N>P>M>G>L: the L mRNA is less transcribed than the other, with a coverage often below 750, which was the threshold of this type of analysis (Figure S5A). Score A, Score B, and Score C were calculated to identify methylated positions as already described beforehand (34). When a position obtained a Score A ≥ 0.3, a Score B ≥ 1, and a Score C ≥ 0.75 compared to their negative control (mix of *in vitro* transcribed RNAs, mix IVT), the position was considered as ribomethylated (Figure S5B). Eleven methylated sites were identified (Figure 4E). Interestingly, only one position was specifically detected after Tha infection (C_m_5103), three positions were specific to Tha-L_K1830R_ (A_m_2442, A_m_2448, and Gm2830), and five positions were specific to Tha-L_T1681S_ (U_m_691, A_m_1750, A_m_2107, A_m_2136, and A_m_2547). Some positions are common to several viruses: U_m_2100 was found at Tha and Tha-L_K1830R,_ and A_m_2481 at Tha and Tha-L_T1681S._ The RT-Low dNTP (RTLP) approach was used to validate the 2’O methylated candidates (Figure S5C). The RTLP technique was validated first on a 100 nt long synthetic RNA containing a 2’O-methylated site that mimics the sequence surrounding position A_m_1909 (Figure S5D). Then positions U_m_2100 and A_m_2547 (Figure S5 E-F) were tested using the same approach; however, no significant results were obtained, which did not allow us to confirm the presence of 2’O methylation on these residues using this ortholog technique.

### Modulation of m^6^A methylation in internal positions of viral mRNA

We next investigated the presence of m^6^A methylation in viral mRNA. To do so, purified mRNA from cells infected with Tha, Tha-L_K1830R_, and Tha-L_T1681S_ were sequenced onto RNA flowcells (Nanopore). Direct RNA-Sequencing determines the RNA sequence by measuring the electrical current as RNA molecules pass through nanopores. Since modified nucleotides produce distinct signal patterns compared to their unmodified counterparts, it becomes possible to computationally detect modifications on a per-molecule basis, using either supervised learning models or comparative analysis methods. Triplicates were sequenced in parallel to a negative control (mix IVT).

Reads were aligned to the viral genome using dorado_align with default parameters. The yield of flowcells varies between 500,000 and 7,000,000 reads. As shown in Figure S6, the proportion of viral reads varied between flowcells.

Methylation rates were computed using modkit pileup for each replicate, and differential modification analysis was performed using modkit dmr pair for the three pairwise comparisons between samples (Tha-L_K1830R_ vs Tha, Tha vs Tha-L_T1681S_, and Tha-L_K1830R_ vs Tha-L_T1681S_). To identify confidently modified positions, we applied the following criteria: p-value < 0.05, a high confidence score, and an effect size > 30%. Only one position met all these criteria: position 68, located in the promoter region of the N gene. This position was significantly more methylated in Tha-L_K1830R_ mRNAs than in Tha and Tha-L_T1681S_ mRNAs (Fig. 4F). In contrast, no significant difference was observed between Tha vs Tha-L_T1681S_, consistent with their overall similarity in the previous experiments.

As described in the Materials and Methods section, a mix of non-modified *in vitro* transcripts (IVT controls) was sequenced (one replicate) and compared to three samples. All conditions exhibited modification at position 68, but with varying degrees: Tha-L_K1830R_ showed the highest methylation level (89%), followed by Tha-L_T1681S_ (50%) and Tha (42%). These levels were determined by comparing each sample to the IVT control, assumed to be unmodified, allowing for the estimation of absolute methylation levels rather than relative differences. These results demonstrate that while position 68 is consistently modified across all samples, the Tha-L_K1830R_ condition exhibits a markedly higher methylation level at this site, suggesting condition-specific regulation of methylation at this site. In addition to position 68, the comparison with the unmodified IVT control revealed consistent methylation at position 70 across all three conditions (Figure 4F). Among the samples, Tha showed the highest methylation level at this site (40%), followed by Tha-L_K1830R_ (32%) and Tha-L_T1681S_ (29%). The relatively small differences in methylation levels between the three different viral mutants prevented this position from reaching statistical significance in the differential analysis performed by modkit dmr pair.

Thus, our results highlight that two positions are marked by m^6^A in the 5’UTR region of the N mRNA: a first one at position 68, which is found with a higher frequency for the Tha-L_K1830R_ mutant, and another one at residue 70, in equivalent proportions between the three viruses.

### Mutation on residue K1830R of the L protein attenuates RABV virulence *in vivo*

The pathogenicity of recombinant Tha-L_K1830R_ was compared to that of Tha in 8-week-old C57BL/6J mice infected with 100 focus-forming units (FFU) by intracranial route. Tha infection caused severe neurological symptoms in all mice starting at 7 days post-infection (dpi) with an intense loss of weight, and all mice succumbed to infection or reached an unresponsive endpoint 10 dpi (Figure 5A-B). Intriguingly, with Tha-L_K1830R_ virus infection, symptom onset and endpoint occurred 10–15 days later, and weight loss was also delayed depending on the appearance of clinical signs (Figure 5A-B). Four animals infected by Tha-L_K1830R_ mice survived >32 dpi.

**Figure 5.**
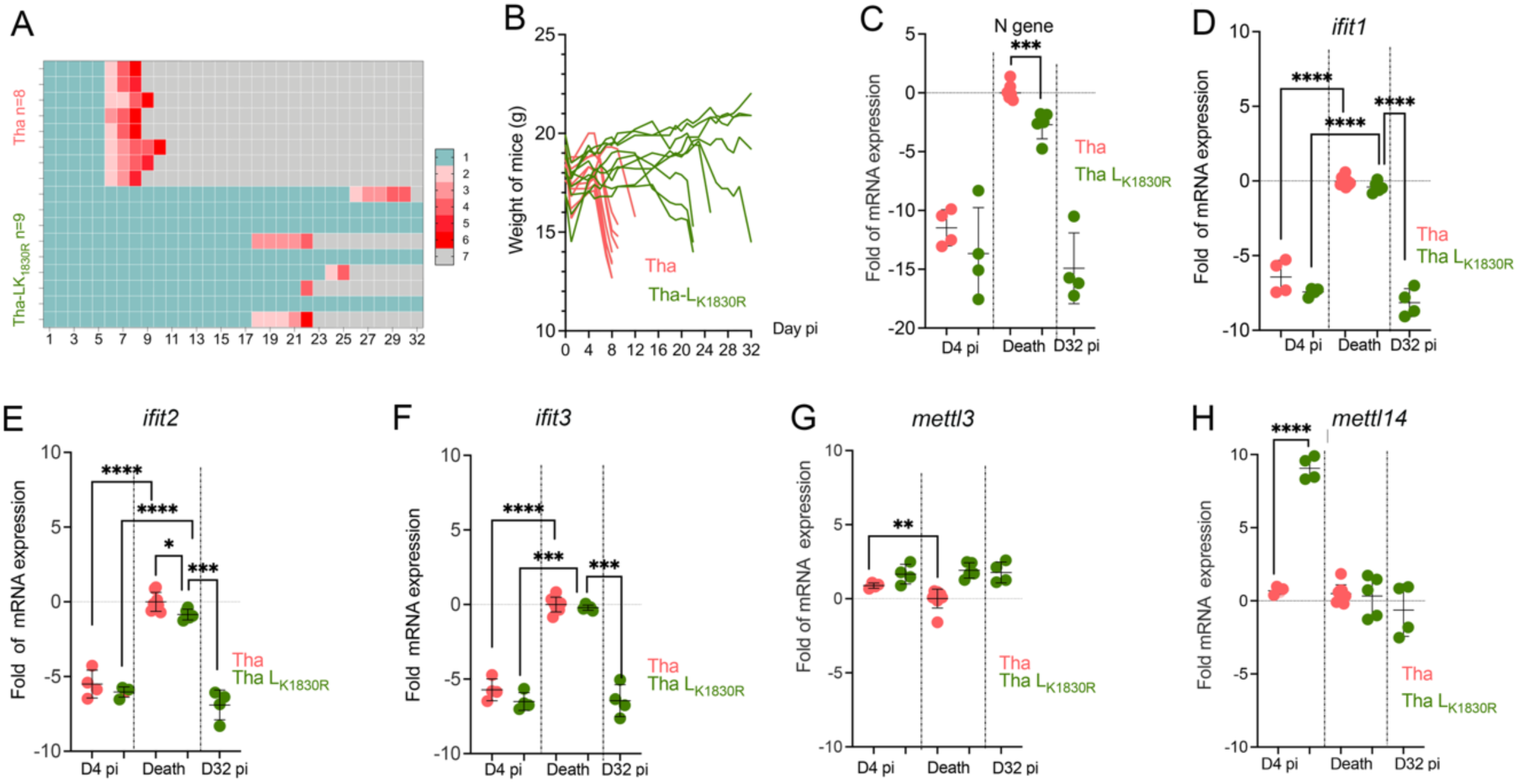
Tha-L_K1830R_ is less pathogenic for mice. Groups of 12 and 13 C57BL/6J mice were respectively infected with Tha and Tha-L_K1830R_ by intracerebral injection. At day 4 post-infection, before the appearance of clinical signs, 4 animals of each group were sacrificed. The brain of the other animal are recovered just before the endpoint or at day 32, when the animal still survives. **(A)** Follow-up of the clinical signs of mice during infection with Tha and Tha-L_K1830R_. Heat maps were established based on a progressive 0–7 clinical score scale (0: no apparent changes; 1: ruffled fur; 2: slow movement, hind limb ataxia; 3: apathy; 4: monoplegia; 5: hind limb paralysis, tremors; 6: paralysis, conjunctivitis/keratitis, urine staining of the haircoat of the perineum; 7: death). Each line represents one animal throughout time. **(B)** Body weight progression of mice during infection with Tha and Tha-L_K1830R_. Mice were weighed daily during the treatment administration and then twice a week up to 32 days post-infection. Each curve represents the body weight of each animal individually. **(C)** Viral nucleoprotein gene expression. RNA was extracted from the brains of the dead animals. Animals sacrificed at D4 (n=4) are indicated by light color. Animals infected with Tha-L_K1830R_ sacrificed at day 32 are indicated by triangles. The fold mRNA expression is calculated relatively to the Tha-infected animals (death at 8.5 days post-infection). Significant results are calculated using an unpaired t-test with Welch’s correction (***: p<0.001). **(D-H)** Gene expression values of *IFIT1*, *IFIT2*, *IFIT3*, *METTL3,* and *METTL14,* respectively. Animals were divided into three groups: animals sacrificed at D4 (n=4), died animals infected with Tha and Tha-L_K1830R_, and animals infected with Tha-L_K1830R_ sacrificed at day 32. The fold mRNA expression is calculated relatively to the Tha-infected animals (death at 8.5 days post-infection). The results represent the mean ± SEM for each group of animals (n=8 or n=9). Significant results are calculated using unpaired t-test with Welch’s correction (****: p<0.0001).

Total RNAs were extracted from the brains of mice infected by Tha or Tha-L_K1830R_, at D4 dpi, at the experimental endpoint, and at D32 pi for the surviving animal. Notably, Tha-infected animals had a significantly higher (***, p<0.001) viral load in the brain than animals infected with Tha-L_K1830R_ (Figure 5C). All these data indicate that viral propagation *in vivo* was impaired by the K1830R mutation in the MTase domain of the L protein.

To determine if those perturbations in infectivity were due to an efficient host response, the expression of mRNAs encoding *Ifit1*, *Ifit2*, and *Ifit3* was measured in the mice’s brains at 4 dpi (Figure 5D-F), at the endpoint, and at 32 dpi for the surviving animal. Both groups of infected animals presented similar *Ifit* expression patterns. *Ifit2* and *Ifit3* mRNA expression increase between 4 dpi and death, whereas *Ifit1* shows a decreased expression level. Conversely, no increase in *Ifit* expression was observed in the brain of the surviving animals (Figure 5D-F). We previously showed that *METTL3* and *METTL14* seemed to be regulated upon *in vitro* infection with Tha-L_K1830R_ (Figure 2I-J). We observed *in vivo* that *Mettl14* mRNA expression was significantly upregulated (****, p<0.0001) upon Tha-L_K1830R_ compared to Tha in the early phase of infection (Figure 5H), with a much higher regulation than observed *in vitro* (Figure 2J). In contrast, *Mettl3* mRNA expression (Figure 5G) was only weakly down-regulated upon Tha infection (Endpoint versus D4; **, p<0,01), conversely to Tha-L_K1830R,_ which was not modulated upon infection.

## Discussion

The innate immune system serves as the primary defence against viral infections, relying on the recognition of foreign nucleic acids and the rapid induction of IFN pathways. Although, it is well established that RABV proteins enable viral evasion of the innate antiviral response, few studies have explored how epigenetic modifications on viral RNAs occur during RABV infection in this process.

Many viruses that replicate in the cytoplasm, such as RABV, encode their own 2’O MTase to convert cap-0 into cap1 structure on viral transcripts, enabling the virus to evade the innate immune response. Indeed, human (HCoV-229E) and mouse (MHV-A59) coronaviruses, Japanese encephalitis virus (JEV), and SARS-coronaviruses mutants lacking 2’O MTase activity induce higher expression of type I IFN and are more sensitive to type I IFN response (3, 16, 35, 36), triggering a signal transduction cascade that initiates an antiviral response and expression of ISGs. Similarly, lyssaviruses also rely on the L protein, which plays a crucial role in genome replication and RNA transcript maturation via the addition of a cap structure at their 5’ end and its subsequent methylation by the MTase domain of the L protein. It is already known that RABV viral mRNA is 2’O-methylated on its cap structure (37).

Our study provides insights into the epitranscriptomic changes observed upon RABV infection, with particular emphasis on the role of the viral MTase and its specific residues facilitating these changes (Figure 6).

**Figure 6.**
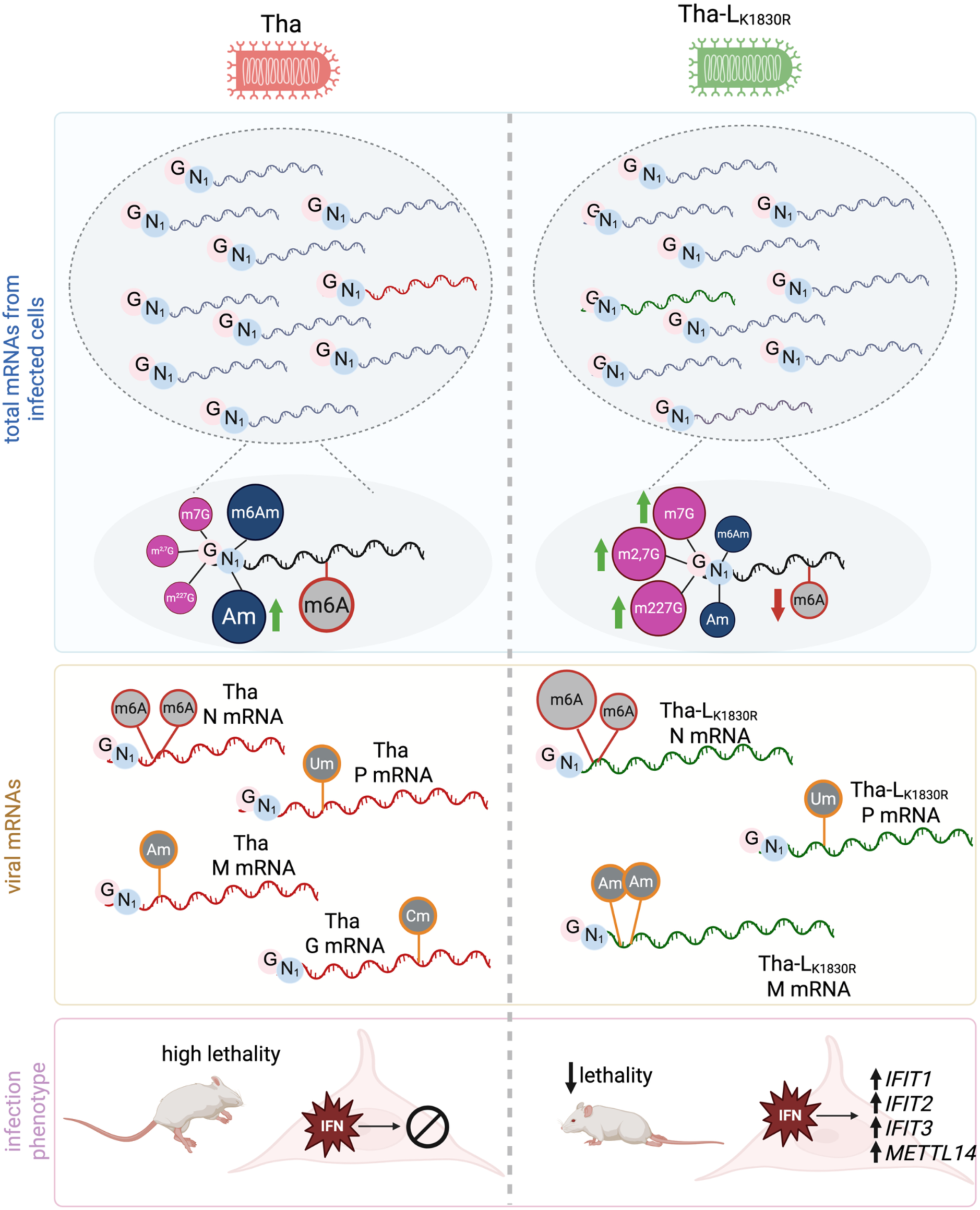
RABV-induced methylation landscapes and associated phenotypes. Tha infection and Tha-L_K1830R_ induce different methylations observed in total RNAs extracted from infected cells (blue panel) and in viral RNAs (light brown panel). It results in different infection phenotypes (pink panel). Tha-infected cells RNAs are more A_m_ methylated on the first nucleotide (included in the cap) than non-infected cells RNAs. Tha-L_K1830R_ -infected cells RNAs are more m^7^G, m^2,7^G, m^2,2,7^G methylated on the cap’s Guanosine and present less internal m^6^A m methylations than non-infected cells RNAs. Tha-L_K1830R_ infection induces higher m^7^G, m^2,7^G, m^2,2,7^G methylation rates on the cap’s Guanosine than Tha infection, and Tha infection induces higher Am and m6Am methylation rate on the first nucleotide, and higher internal m^6^A methylation rates than Tha-L_K1830R_. Green arrows indicate a higher methylation rate that the one observed in non-infected cells. Red arrows indicate a lower methylation rate that the one observed in non-infected cells. Size of the methylation size of the methylation is proportional to the methylation rate and enable comparison of methylation rates between Tha infected cells RNAs/viral RNAs and Tha-L_K1830R_ infected cells RNA/ viral RNA. G: capped guanosine; N1: first nucleotide; Grey RNA: cellular mRNA; Red RNA: Tha viral mRNA; Green RNA: Tha-L_K1830R_ viral mRNA.

To determine whether internal 2’O methylations are present specifically in the RABV RNAs, a high-throughput sequencing approach (RiboMethSequencing) was developed in our laboratory, and we elucidated for the first time the specific internal 2’O-methylations modifications of RABV mRNAs (Figure 4E). By using RiboMethSeq, we detected the presence of a small subset of internal 2’O-methylations in RABV mRNAs; most of the internal methylations were on A residues, and the N_m_ were mainly detected in Phosphoprotein, Matrix, and Glycoprotein mRNAs after infection with Tha virus. Unfortunately, none of those positions were confirmed using the orthologue method (RTLP approach), possibly because of the low rate of mRNA methylation (<20%), which is the threshold of the RTLP approach. This could be because these methods only allow the identification of N_m_ if they are systematically present on a specific residue. If the methylations are deposited randomly, it is impossible to conclude from those studies. Recently, it was demonstrated that Ebola virus MTase (38), as well as Dengue and Zika virus MTases (8, 39), also induce specific internal adenosine 2’O-methylations *in vitro*. The role of such methylations is still under debate, but in the case of HIV-1 viral RNA (13), internal methylations were found to limit the detection by the RLR MDA5 sensors and, in turn, reduce type-1 interferon secretion. Thus, if such methylations are present within viral mRNAs, it might allow RABV to evade these pattern recognition receptors.

We also looked at internal m^6^A methylation using nanopore sequencing. It allowed us to identify two m^6^A modifications in the 5’UTR region of the N mRNA (residues 68 and 70 on the Tha sequence) present in wild-type Tha virus (Figure 4F). The mRNAs containing *N*^6^-methyladenosine in their 5′ UTR can be translated in a cap-independent manner through eIF3 instead of the canonical pathway involving eIF4E (40). In this case, 5′ UTR m^6^A bypasses 5′ cap-binding proteins to promote translation under stress conditions. This regulation of N protein translation could give the virus an advantage in producing enough N protein to efficiently encapsulate viral RNA and escape host recognition to compensate for interferon susceptibility. A further explanation could be that N mRNAs are the most prevalent viral mRNAs during viral transcription, which increases the probability of finding these methylations, which are, however, only slightly represented (approximately 30% compared to IVTs).

Our data also confirmed that the first cap structures of mRNA from infected cells were 2’O methylated and showed that these first nucleotides were also N6-2’O methylated: in the case of the wild-type virus, Am methylations are overexpressed compared to healthy cells (Figures 4B). However, the different sequencing methodologies revealed that the mRNAs purified from infected cells contained only between 2-20% of viral mRNAs (41), indicating that modification of methylations should also affect cellular RNAs. This could be a direct effect of the viral MTase on cellular mRNA, but it could also result from indirect effects of the virus or from the cellular response to infection.

To have a better understanding of the mechanisms underlying internal and cap 2’O and m6A methylations, we looked closer into the RABV MTase domain. We generated site-specific mutants inside the K-D-K-E motif. The effect of each mutation introduced in the catalytic tetrad residues motif was analyzed first by a minireplicon using the CAT assay. Those mutations were deleterious for the replication activity, and only L_K1830R_ still showed weak replication. Each residue of the K-D-K-E tetrad is known to have an effect on N7 and 2’O MTase activity of the L protein. Mutagenesis studies have previously shown that any mutation in the K-D-K-E catalytic tetrad of the flavivirus MTase almost completely abolishes its 2′O-MTase activity, whereas the N7-MTase activity is likely to mainly depend on the presence of the D and E residues of the catalytic tetrad (42). Interestingly, when mutations were introduced by reverse genetics into the complete RABV genome (our study), mutation on residue D was not stable and reverted quickly on the same position to the wild-type sequence. Mutation on E was very deleterious for the virus (Figure S2), and it was not possible to obtain a viral stock, suggesting that this residue plays a key role in both N7 and 2’O methylation. Conversely, virus bearing mutations K1686R or K1830R were usable, suggesting that at least the N7-MTase activity is conserved. Their growth kinetics were determined on IFN-competent (SK-N-SH) and IFN-non-competent (BSR) cells. Surprisingly, Tha-L_K1830R_ growth was impacted in both IFN-non-competent and IFN-competent cells, whereas Tha-L_K1686R_ was only impaired in SK-H-SH cells.

This suggests that the impact of these mutations is not exactly the same on the 2′-O position, as K1830R displays a more pronounced replication phenotype. As control, we introduced the mutation T1681S, located at the beginning of the MTase domain, and observed no significant difference in growth between Tha-L_T1681S_ and Tha viruses. This amino acid, localized at position 1670 of hMPV L protein specifically was previously demonstrated to limit its 2’O-MTase activity (23), but in the case of RABV, no impact was shown in viral growth or IFN pathway activation

The potential antiviral response induced by catalytic mutants was assessed in STING-37 cells expressing luciferase (Figure 2E) under the control of the IFN-stimulated response element (ISRE). As expected, the Tha-L_K1830R_ induced a weak but significant increase in luciferase activity on STING37 compared to Tha, and even more after IFN-α treatment, showing higher activation of ISRE and then suggesting the detection of mis-capped viral RNA by RIG-I. Conversely, the virus mutated at the T1681 residue is still able to totally restrict the host response, probably through the action of its P and M proteins (26, 27, 29, 30). Indeed, activation of the ISRE promoter in Tha-L_K1830R_ cells is not sufficient to induce *IFIT* expression, as seen in Figure 2F-H without IFN-α stimulation during infection of the SK-N-SH. On the other hand, when IFN-α is added exogenously, Tha-L_K1830R_ virus stimulates more efficiently the ISRE promoter at an efficient level that induces an increased gene expression of *IFIT1*>*IFIT3*>*IFIT2* genes which is not observed using Tha virus (Figure 2F-H). The IFIT family plays an important role in the destruction of non-self RNA. In collaboration with other ISGs proteins, they can restrict viral replication. For example, viral RNA carrying a 5’ppp extremity can be recognized by IFIT1, followed by sequestration with the IFIT complex, which contains IFIT1, IFIT2, and IFIT3. Moreover, human IFIT1 has been shown to inhibit replication of viruses lacking N1 ribose 2′-O methylation, such as HCoV 229E bearing a D129A (DA) mutation in its viral 2′-O MTase gene (43), and interaction between IFIT1 and IFIT3 is important for restricting infection of viruses lacking *2′-O* methylation in their RNA cap structures. Thus, the presence of IFIT1 requires that viruses replicating in the cytoplasm maintain mechanisms to avoid IFIT1’s restrictive effects.

In the case of cells infected with the mutant virus Tha-L_K1830R,_ impaired in its 2’O MTase activity, 2’O methylation on the first nucleotide of the cap structure of mRNA is decreased after infection compared to wild-type Tha virus. This is correlated with *IFIT1*, *IFIT2,* and *IFIT3* mRNA upregulation compared to Tha. Therefore, our data confirm that 2′-O-methylation of the first transcribed nucleotide could serve as a “self” signal that helps cellular RNAs evade detection (3, 44). Conversely, methylations on the guanosine of the mRNA cap structure (m^7^G, m^2,7^G, and m^2,2,7^G) are significantly increased (Figure 4A, S4A, B). Our data also revealed differences in internal methylation of RNAs from infected cells. Upon Tha-L_K1830R_ infection, both m^6^A and A_m_ are decreased in comparison to mRNA extracted from Tha-infected cells (Figure 4C).

Nanopore sequencing also showed that the two m^6^A modifications in the 5’UTR region of the N mRNA (residues 68 and 70 of Tha complete sequence, corresponding to residues 10 and 12 in N mRNA) were also present in Tha-L_K1830R_ mRNA but in different proportions. Position 10 is highly methylated in the case of the Tha-L_K1830R_ virus (approximately 89%), whereas it is only present in 30% of cases for the other two mutant viruses. The m^6^A methylations are catalyzed by the m^6^A MTase (writer) complex, which includes MTase 3 (METTL3) and MTase 14 (METTL14), and both mRNAs were upregulated upon Tha-L_K1830R_ infection *in vitro* but also *in vivo* (Figure 2I-J, Figure 5H-I). It suggests that the L-_K1830R_ mutation changes the global methylation landscape of the mRNA pool of infected cells and, in turn, perturbs the host response. All these modifications are correlated with a higher sensitivity to IFN-α *in vitro*, and less pathogenicity *in vivo*, as mice infected with intracerebral injection of Tha-L_K1830R_ virus show higher survival and fewer symptoms than mice infected with Tha (Figure 4A-B).

In conclusion, the mutation K1830R in the K-D-K-E tetrad of the Tha MTase domain of the L protein induces numerous changes at the gene expression level, the methylation landscape for both virus and host mRNAs, and leads to a less pathogenic phenotype *in vivo* compared to the wildtype strain (Figure 6). Altogether, these results suggest a regulatory effect of the viral MTase and that cellular MTases participate in viral RNA methylation and may then play a role in the control of RABV infection. Although the exact role of these epitranscriptomic modifications of viral and cellular mRNA is not yet fully understood, some of these methylations, like cap-2’O methylations and internal m6A methylations, appear to have a proviral effect and increase the viral propagation by allowing RABV wildtype strain Tha to efficiently evade the host antiviral response.

## Materials and methods

### Cells and viruses

BSR cells were used for viral titration and production of the viruses. SK-N-SH cells (ATCC® HTB-11™) were used for viral growth studies, gene expression, RiboMethSeq and nanopore experiments. Human epithelial kidney cells stably transfected with an ISRE-luciferase reporter gene-37 (STING37) were kindly provided by Marianne Lucas-Hourani (45) (Unité de Génomique Virale et Vaccination, Virology Department, Institut Pasteur). All cells were cultured at 37 °C, 5% CO_2_, DMEM supplemented with 10% calf serum. BSR-T7 cells provided to us by K.K. Conzelmann (Max von Pettenkofer Institute and Gene Center, Munnich) were cultured in Glasgow medium supplemented with 10% calf serum, tryptose phosphate, non-essential amino acids, and geneticin (46).

Six recombinant viruses derived from rabies strain 8743THA (EVAg collection, Ref-SKU: 014V-02106), a street-dog isolate from Thailand, were used: wild-type virus (Tha) and five L-protein mutants (Tha-L_K1681R_, Tha-L_D1798N_, Tha-L_K1830R,_ and Tha-L_E1868N_, harbouring mutations in the K-D-K-E motif and Tha-L_T1681S_, mutated on a conserved residue upstream the K-D-K-E motif). The different mutations were introduced into the L sequence on the plasmid encoding the complete genome of the Tha using the Phusion Site-Directed Mutagenesis Kit (Cat.#F541, Thermo Scientific) according to the manufacturer’s instructions. All mutagenesis primers are listed in Supplementary Table 1.

### Reverse genetics

Different recombinant viruses were produced by reverse genetic as already described (Ben Khalifa, Luco et al. 2016) by transfection into BSR-T7 cells of the complete genome (2.5 μg), and plasmids N-pTIT (2.5 μg), P-pTIT (1.25 μg) and L-pTIT (1.25 μg). Briefly, 6 days after transfection, supernatants were passaged onto BSR cells and incubated for 5 days. The supernatant was harvested and titrated on BSR cells.

### Infection and viral growth curves

For all experimental conditions, BSR and SK-N-SH cells were infected with Tha and five mutants at a multiplicity of infection (MOI) of 0.5 Foci Forming nit (FFU) /cell. One million cells were seeded in 6-well plates 24h before infection. For infection, supernatant was removed, and the virus/mutants were added by adding 0.5 mL of serum-free medium. After 1 hour of incubation, the medium was removed and replaced with 1 mL of medium supplemented with 2% of SVF, and the plates were incubated for 48h.

For growth curves, 2 × 10^6^ cells (BSR or SK-N-SH) were infected under agitation at 37 °C with 5 % CO_2_. After 1h incubation, the cells were centrifuged at 800 g for 5 minutes, and the pellet was resuspended in complete medium. The cells were then seeded in a T25 flask and incubated at 37 °C with 5 % CO_2_. After 24, 48, and 72 h of incubation, 0.5 mL of supernatant was aliquoted for titration, and 0.5 mL of fresh medium was added to each flask.

### Minireplicon assay

Minireplicon assay was carried out as already described (31). Briefly, BHK-T7 cells were transfected in 6-well plates with either 1 μg of pSDI-THA-CAT (−), 10 ng pCMV-RL, 1 μg of N-pTIT, 0.5 μg of P-pTIT, and 0.5 μg of L-pTIT using 6 μL of lipofectamine 2000 (Cat.#11668019, Invitrogen). For the negative control, one of the pTIT vectors was omitted. Forty-eight hours post-transfection, cells were harvested in passive lysis buffer (Cat.#11363727001, Roche), and lysates were subjected to reporter CAT assays using the ELISA CAT assay (Cat.#11363727001, Roche) according to the manufacturer’s instructions. Each lysate was tested undiluted and on three serial 1:10 dilutions. Luciferase activity was measured by using the Renilla-Glo assay (Cat.#E2710, Promega) according to the manufacturer’s instructions on undiluted sample lysates.

### Cell staining and image acquisition on the Opera Phenix High Content Screening System

SK-N-SH cells were seeded at a density of 1.8x10^4^ cells/cm^2^ in 96-wells plates (#655090, Greiner Bio-One). Twenty-four hours after seeding, cells were infected with Tha, Tha-L_K1830R,_ and Tha-L_T1681S_ at a MOI of 0.5. Cells were incubated with viral suspension at 37°C and 5% CO_2_. After two hours, the viral suspension was removed, and 100 µl DMEM supplemented with 10% FCS were added. Cells were fixed at 24- and 72-hours post-infection. In detail, cells were fixed using 4% PFA (Alfa Aesar, J61984) for 15 minutes at room temperature, washed with PBS (10010023, Thermo Scientific), and permeabilized using 0.5% Triton X-100 (648463, Millipore) for 10 minutes. Cells were stained for 30 minutes with a FITC-labelled monoclonal antibody directed against the RABV nucleoprotein (Sigma-Aldrich, 5100) at 37°C. Subsequently, cells were stained for 5 minutes with Hoechst 33342 (Thermo Fisher, H1399) according to the manufacturer’s instructions. Images were acquired via the Opera Phenix High Content Screening System (Perkin Elmer) using channels 480/500-550 (Nucleoprotein, 200 msec, 50% intensity) and 375/435-480 (Hoechst, 50 msec, 100% intensity). To measure relative fluorescence levels, we used the software Columbus (Perkin Elmer), which automatically detects nuclei and the cellular cytoplasm. Cytoplasmic intensity thresholds were calculated based on the autofluorescence level of non-infected SK-N-SH.

### RNA extraction and RT-qPCR

Total RNA was extracted from *in vitro*-cultured cells using RNeasy mini kit (Qiagen, Hilden, Germany). Total mouse-brain RNA was isolated with TRIzol. RNA was reverse transcribed to first-strand cDNA using the SuperScript™ IV VILO™ Master Mix (11766050, Invitrogen). qPCR was performed in a final volume of 10 μL per reaction in 384-well PCR plates using a thermocycler (QuantStudio 6 Flex, Applied Biosystems) and its related software (QuantStudio Real-time PCR System, v.1.2, Applied Biosystems). Briefly, 2.5 μL of cDNA (12.5 ng) were added to 5 μL of Taqman Fast Advanced master mix (444457, Applied Biosystems) and 2.5 μL of nuclease-free water containing primer pairs and probe (Supplementary Table 1). The amplification conditions were as follows: 95 °C for 20 s, and 45 cycles of 95 °C for 1 s and 60 °C for 20 s. The *gapdh* and the *hprt* (hypoxanthine phosphoribosyl-transferase) genes were used as reference. Variations in gene expression were calculated as the n-fold change in expression from the infected cells or animals compared with the mock-infected group using the 2^-ΔΔCt^ method (47).

### mRNA purification using Dynabeads® mRNA DIRECT™ Kit

Four wells of a 6-well plate were used for each viral infection condition. After 48 hours of infection, the medium was removed, the cells were washed with PBS, and then lysed with 250 µL/well of Lysis/Binding Buffer for 30 minutes at 4°C. Cell lysates were collected, and each well was rinsed with 50 μl of Lysis/Binding Buffer to obtain a final volume of 300 μl. For every condition, two wells were pooled together. Lysates were forced 3-5 times through a 21-gauge needle (307732, BD Emerald) using a 1-2 ml syringe to shear the DNA to reduce the viscosity. The purification on beads was performed according to the manufacturer’s instructions with minor modifications: Cell lysates were incubated on 50 µL of pre-washed oligo d(T) beads, and serial washings were performed with buffer A and B, and mRNA was eluted from the beads by adding 20 μl of RNAse-free water and incubating them at 65°C for 2 minutes. To increase mRNA purification, another purification was performed subsequently. The previous beads were washed two further times in Washing Buffer B, and then the purified mRNA was diluted in 80 μl of Lysis/Binding Buffer. Serial washings are performed with buffer A and B, and a final elution is performed as described above. The quality and the concentration of the mRNA are controlled with the Bioanalyzer RNA nano kit (5067-1511, Agilent) and the Qubit RNA IQ Assay (Invitrogen), respectively.

### Mass spectrometry

350 ng of mRNA purified from infected cells were digested by 25 U of RppH (NEB, #M0356S) for 2 h at 37°C. Decapped mRNAs were then digested by nuclease P1 (NEB, #M0660) for 2h at 37°C. Nucleotides were dephosphorylated for 2h at 37°C by 5 U of alkaline phosphatase (rSAP, NEB#M0371)

After digestion, the samples were filtered with 0.22µm filters (Millex®-GV, Millipore, SLGVR04NL). 5µL of each sample was injected in triplicate in a LC-MS/MS. Nucleosides were separated using Nexera LC-40 systems (Shimadzu) and Synergi™ Fusion-RP C18 column (4µm particle size, 250mm x 2mm, 80Å) (Phenomenex, 00G-4424-B0). Mobile phase is composed of 5mM ammonium acetate adjusted to pH 5.3 with acetic acid (solvent A) and pure acetonitrile (solvent B). The 30-minute elution gradient started with 100% phase A, followed by a linear gradient until reaching 8% solvent B at 13 min. Solvent B was further increased to 40% over the next 10 minutes. After 2 minutes, solvent B was decreased back to 0% at 25.5 minutes. Initial conditions were recovered by rinsing the LC system with 100% solvent A for an additional 4.5 minutes. The flow rate was set to 0.4 mL/min and the column temperature to 35 °C. Analysis was performed using a triple quadrupole 8060NX (Shimadzu corporation, Kyoto, Japan) in positive ion mode. The ESI source settings and the multiple reaction monitoring (MRM) transitions are described in Supplementary Methods. Peak areas were measured using Skyline 21.2 software (MacCoss Lab Software; https://skyline.ms), and the ratio of modified nucleosides to non-modified nucleosides was calculated.

### *In vitro* RABV RNA transcription

Plasmids P2RZ-N, P2RZ-P, P2RZ-M, P2RZ-G and P2RZ-L were generated using the inFusion cloning kit (Takara). DNA fragments corresponding to nucleoprotein (N), phosphoprotein (P), Matrix, glycoprotein (G), and Large protein (L) of rabies virus were amplified by PCR using specific primers (Table 1).

The five different plasmids were linearized using the EcoRI restriction enzyme. P2RZ-M and P2RZ-L were linearized by *HindIII* (Thermo Fisher Scientific). T7 transcription reactions were carried out with a T7 RiboMAX Express Large Scale RNA Production System (Promega). The DNA strand was digested using RQ1 DNase (<u>Promega</u>). RNAs were purified using ethanol precipitation and glycogen and analyzed with Bioanalyzer RNA nano kit (Agilent).

### RiboMethsequencing library construction

Total RNAs were recovered from cells infected with the three recombinant viruses: Tha, Tha-L_K1830R,_ and Tha-L_T1681S,_ and then viral mRNAs were extracted using Dynabeads mRNA direct kit as already described. Two hundred ng of viral mRNAs extracted from viral particles or an equal amount of a mix of *in vitro* transcribed RNAs (mix IVT) used as negative control, were submitted to alkaline fragmentation in 50mM bicarbonate buffer pH 9 for 8 min at 95°C. The reaction was stopped by ethanol precipitation using 3M Na-OAc, pH 5.2, and glycogen. After centrifugation, the pellet was washed twice with ethanol 70% and resuspended in nuclease-free water. The RNA fragments were converted to a library using NEBNext Small RNA Library Prep Set according to the RiboMethSeq protocol described in Birkedal et al. (48). The obtained oriented libraries were controlled on Bioanalyzer DNA1000 Chips (Agilent, # 5067-1504) and quantified by spectrofluorimetry (Quant-iT™ High-Sensitivity DNA Assay Kit, #Q33120, Invitrogen). Sequencing was performed on the Illumina HiSeq 2500 platform to generate single-end 65 bp reads bearing strand specificity.

### RiboMethSeq bioinformatic analysis

Sequencing reads were first processed to remove adapter contamination and low-quality sequences using Cutadapt v1.11 (49), retaining only reads of at least 25 nucleotides in length. Cleaned reads were aligned to the rabies virus Tha reference genome (Lyssavirus rabies isolate 8743THA, accession number EU293121.1) using Bowtie v1.2.2 (50) with the -m 1 option to keep only uniquely mapping reads.

The 5′ and 3′ end positions of aligned reads were extracted and used to compute site-specific cleavage profiles. RiboMethScores were then calculated following the method described by Birkedal et al. (48) using a custom Python script. The scoring was performed using a ±2 nucleotide window around each position, yielding three metrics: Score A, Score B, and Score C. To identify methylated positions, we applied the filtering thresholds adapted from Pichot et al. (51): Score A ≥ 0.3, Score B ≥ 1, and Score C ≥ 0.75. Comparisons of RiboMethScores and coverage between viral strains enabled the identification of differentially methylated sites. Positions are marked as ribomethylated when they passed these filters and the negative control (mixTIV) did not. 5’ and 3’-ends coverage were normalized with DESeq2, ribomethylated positions coverages and scores are plotted using ggplot2 under R version 4.0.3 using ggplot2.

### RTLP and validation of methylated candidates

For the validation of 2’O-methylated candidates, extracted viral mRNAs were used for RT-PCR in 25 μl reaction mixture containing 100 ng of mRNA, 10 μM of specific RT primers, and a low (0.5 μM) or high (1 mM) concentration of dNTPs. The Primer/RNA mixture was denatured at 70°C for 5 min, then chilled on ice. Following an initial annealing step at 42^°^C for 10 min, 200 U of M-MLV reverse transcriptase (Promega) and 0.5 U RNasin Ribonuclease Inhibitor (Promega) were added. The reaction was incubated at 42^°^ C for 1 h and then heated at 75^°^ C for 15 min to deactivate the reverse transcriptase. A mix of *in vitro* transcribed RNAs was used as a negative control.

### Direct RNA Nanopore sequencing

The previously purified mRNAs were concentrated using Monarch® Spin RNA Cleanup Kits (ref T2030S) to be able to recover around 300 ng of poly(A) tailed RNA. RNA libraries were prepared using the Direct RNA sequencing kit (Nanopore, SQK-RNA004) according to the manufacturer’s protocol. Library preparation began with the ligation of a specific adapter to the 3’ end of the RNA to allow reverse transcription initiation. This step was followed by the synthesis of a complementary strand of cDNA using reverse transcriptase. Although this strand is not sequenced, its presence stabilizes the RNA molecule during the sequencing process. Once the RNA-cDNA hybrid was formed, nanopore-specific sequencing adapters were ligated to the ends of the molecule to allow its recognition and translocation into the pore. Each enzymatic reaction was followed by magnetic bead purification to remove excess reagents and unbound adapters. The resulting library quantity was then quantified by fluorimetry using the Qubit 4 and the Qubit dsDNA HS Assay Kit (Invitrogen, ThermoFisher Scientific, #Q32851). Libraries were loaded onto RNA flow cells (FLO-MIN004RA), which had previously been checked (Flowcell Check). 72h sequencing was performed on a GridION (Oxford Nanopore Technologies). MinKNOW software was used for data acquisition and real-time basecalling. through the nanopore into the corresponding base sequence of the strand.

### Direct RNA Nanopore bioinformatic analysis

Basecalled reads passing a quality score >7 were processed using Dorado v1.0.0 https://github.com/nanoporetech/dorado) with the high-accuracy model rna004_130bps_hac@v5.1.0.

RNA modifications were detected using the inosine_m6A_2OmeA model to identify m6A sites across all samples. Basecalled reads were aligned to both the human reference genome (hg38, https://www.ncbi.nlm.nih.gov/datasets/genome/GCF_000001405.40/) and the rabies virus genome using dorado_align. Methylation rates were then computed for each replicate using Modkit pileup v0.5.0 (https://github.com/nanoporetech/modkit), with the modification detection threshold determined automatically by the software. Differential modification analysis between two conditions (three replicates per condition) was performed using modkit dmr pair. M6A sites with a balanced p-value < 0.05 were kept.

### *In vivo* experiments

All animal experiments were performed according to the French legislation and in compliance with the European Communities Council Directives (2010/63/UE, French Law, 2013–118, February 6, 2013) and according to the regulations of the Pasteur Institute Animal Care Committees. The Animal Experimentation Ethics Committee (CETEA 89) of the Institut Pasteur approved this study (160111; APAFIS#l5773-2018062910157376 v5). All animals were handled in strict accordance with good animal practice.

Eight-week-old female C57BL/6J mice were purchased from Charles River Laboratories and handled under specific pathogen-free conditions, according to the institutional guidelines of the Central Animal Facility at the Institut Pasteur, with *ad libitum* access to water and food. Before any manipulation (surgery or infection), animals underwent an acclimation period of 1 week. Mice were anesthetized intraperitoneally with 100 mg/kg ketamine (Imalgène 1000, Merial) and 10 mg/kg xylazine (Rompun, Bayer), and then were infected with 100 FFU (fluorescent focus units) of the pathogenic Tha-RABV or the recombinant Tha-L_K1830R_ one, in a volume of 30 μl, injected by intracranial route. Thereafter, animals were monitored daily, with body weight and clinical signs noted. The clinical signs were classified in a progressive 0–7 scale (0: no apparent changes; 1: ruffled fur; 2: slow movement, hind limb ataxia; 3: apathy; 4: monoplegia; 5: hind limb paralysis, tremors; 6: paralysis, conjunctivitis/keratitis, urine staining of the haircoat of the perineum; 7: death). In the clinical phases of the disease, when the animals were paralyzed, DietGel Recovery (#72-06-5022, ClearH_2_O) was used as a diet supplement.

### Statistics

Statistical tests were carried out using Prism software (GraphPad Prism, version 10, San Diego, USA), with p < 0.05 considered significant. Non-linear regression analysis, Mann-Whitney tests, and unpaired *t*-test with Welch’s correction were performed.

## Acknowledgment

We acknowledge all the members of the Lyssavirus, epidemiology, and neuropathology team for their help and useful discussions.

We thank the members of the Biomics platform at Institut Pasteur for providing advice for NGS experiments.

This study was supported by ANR-16-CE11-0031-01, the Institut Pasteur, and the CNRS. Rania Ouazahrou, from Biomics Platform, C2RT, Institut Pasteur, Paris, France, was supported by France Génomique (ANR-10-INBS-09) and IBISA.

