## supplementary data.pdf for "RABV L protein plays a role in immune escape through its methyltransferase activity"

| | | |
|---|---|---|
| **Consensus** | ----MIDXXEVYDD-PXDPVEPEXDXXXN-XVVPNILRNSDYNLNSPLLE |  |
| RABV-Tha | ---ML..SG.....-.V..I.S.AEPRG-SPT................I. | 45 |
| ARAV | ----...PL.....-.V......AEFRG.-S.................I. | 44 |
| ABLV | ----...PG.....-.I......PELKT.NA.I...............I. | 45 |
| BBLV | ----.L.PQ.....-.V......T.LKN.-S.................I. | 44 |
| WCBV | ----.LEST.....-.L......Q.WRPE-ASA................S | 44 |
| DUVV | ----.LEST.....-.L......F.LKT.-S................... | 44 |
| TWBLV | ----.LEAT.....-.L......AEFKN.-S.I................. | 44 |
| GBLV | ----...PG.....-.I......V.LRN-NPTI...............I. | 44 |
| EBLV1 | ----...ST.....-.V......A.LRS.-S................... | 44 |
| EBLV2 | ----...PL.....-.V......I.ARS.-S.................I. | 44 |
| IKOV | -----M.WS..I..-.E..FSLDEESCSV-D.P.....T........... | 43 |
| IRKV | ----..EST.....-.V.....DV.IRN.-S................... | 44 |
| KHUV | ----...SL.....-.V..I...V.LKN.-A.................I. | 44 |
| KBLV | ----.MEPP.....-.S......A.IKN.-S.................I. | 44 |
| LBV | ----...SS.....-.I..A...CEWSG.-P.I................. | 44 |
| LLEBV | -----M.WS..T..-.E.LI.Y.P.SVSG-DPP.....T........... | 43 |
| SHIBV | ----..ESS.....-.L..A...SEWSNT-SII................. | 44 |
| MOKV | ----.M.IT.....-.I......GEWNSS-P................... | 44 |
| VSV_NJ | MDFDL.EDSDNWG.DES.FFLRDI--L-SQEDQMSY.NTA........IS | 47 |

| | | |
|---|---|---|
| **Consensus** | DPSXLMLXWLXTGNXPXRLXXTDNXXRSYKVLKXYFKKXDXGSXKXGGXG | |
| RABV-Tha | ..AR...E..K...R.Y.MTL...CS.....F.D....L.L..V.V..TA | 95 |
| ARAV | ...R...E..T...K.N.MTF...SL.......S....I.V..LRV.SS. | 94 |
| ABLV | H..R...E..T...K.M..TL...CI....T..C..R.V.I..V.V..PA | 95 |
| BBLV | ...R...D..T...R.N..NM...AF.......S...RL.V..L.V..V. | 94 |
| WCBV | ...E..IS..QS.QK.N.IPLSE.IR..Q.I..LSL..V.LATVLT..DS | 94 |
| DUVV | ...K...E..M...K.H..NL...LL.......G...RL.T..LRV.SV. | 94 |
| TWBLV | .S.R...E..G...R.N..TP...LL......RN..R.R.V..V.I..P. | 94 |
| GBLV | ..AR...E..TS..K.N.MTL...CL.......G....V.I..L.V..PA | 94 |
| EBLV1 | ...R..IE..T...R.S..NL...LL......RG....L.V..MRA..L. | 94 |
| EBLV2 | ...K...D..I...K.A..NL...SL.......S....L.V..LRV..L. | 94 |
| IKOV | .Q.L..IE..K.A.K.S..RHSEGLL....T..DCL..I.LD.LRY..Q. | 93 |
| IRKV | ...R...D..I...K.C..NL...LF......RS....I.I..L.V.SL. | 94 |
| KHUV | ...Q...D..I...K.N..SPS..LV.......S....V.V..LRI..L. | 94 |
| KBLV | ...K...D..I...R.N..SV...SV.......G....L.V..LRV..P. | 94 |
| LBV | ..TH..IQ..LS..R.L.MTV.E.MV..H....AS..RV.IAVI.I..TS | 94 |
| LLEBV | .SCQ..IE..K...R.S..KSS.GLI..H.A..ECL.RA.LETV.Y..S. | 93 |
| SHIBV | .HAD...Q..SS..R.L.MST...IS.......LS...I.IA.I.F..Q. | 94 |
| MOKV | ..AN..IQ..TS..R.M.MNV.E.TN...R...AF..GI.VATV.I..A. | 94 |
| VSV_NJ | .DIVYIVKRMNHEDV.PIWRSKE-WDSPLDM.RGCQ------AQPMSHQE | 90 |

```
Consensus   AQAMXXLWLHGSHSESTRSRKCLXDLXXFYQKSSPIEKLLN-YTLENRGL
RABV-Tha    ..S.IS...Y.A....N...R.IA..AH..S..........-C..G....   144
ARAV        ....IS..I.......N......T..SN.............-........   143
ABLV        ....TS.....E....N......S..TQ.............-...G....   144
BBLV        ....MS.....A...........S..TH...R.........-........   143
WCBV        SRSVMA..CNK..L......R..NN.SQ..N......AI.K-.S..I...   143
DUVV        S...VT.................AE.SQ..K..A...T...-C.......   143
TWBLV       ..S.IN..I..............TE.SQ..K..........-........   143
GBLV        ....IS.....A....N.....MS..AQ..H.........T-...G....   143
EBLV1       ....IT.................TE.SQ..K..........-........   143
EBLV2       ..S.MS.....A...........S..AL...R.A.......-........   143
IKOV        .KLAFKR.VTHNYV....N..MFM..RE....LE...TIT.-RF..R...   142
IRKV        ..S.TT.......A.........S..SQ..K......I...-........   143
KHUV        ....TS.....A...........S..AQ.............-........   143
KBLV        ....IS.....A...........S..TQ...R.........-........   143
LBV         ....MGT.VR...A..S......A..SS...R.....SI..-...M....   143
LLEBV       .KMVFKK.VTN.YV....N..MFA..QE....VE...AITS-KF..K...   142
SHIBV       ....MNT.VLC..A..S......T..SI...R.I...SI..-...S....   143
MOKV        ....MG..VL..N...S...R..T..SA..RRTM...SI..-H..M....   143
VSV_NJ      MHNWFGT.IQNIQHD.AQGFTF.KEVDK---E.EMTYD.VSTFLKGWV.K   137
```

```
Consensus   XXPXEGVLSSLXKVXYDXAFGRYLGNXYSSYLFFHVIILYMNALDWDEEK
RABV-Tha    RI.P.....C.ERID..K......A.T..........T........E...   194
ARAV        KI.R.......R..N..H........T.A......I..............   193
ABLV        RI.P....LC.K..D..RS.....A.I.......................   194
BBLV        QT.P.......K..D..Q.....M..V...................E...   193
WCBV        QN.RD.I.T..E..N..SS....M..V.A...L...V.............   193
DUVV        RT.P.....C.D..H..Q........T.........MV............   193
TWBLV       NT.S....T..D.IQ..Q........T.........VV............   193
GBLV        RI.S....AC.K..D..K......A.I..........V............   193
EBLV1       KT.T......IN..Q..Q........T.......................   193
EBLV2       AI.TD......K..N..R........L...................E...   193
IKOV        QC.SL.I.E.FKTLSP.S.L...FV.V.A...V....M....S.......   192
IRKV        QT.A.......N..H..Q........T..A....................   193
KHUV        QV.P.......K..S..R......A.T...................E...   193
KBLV        QT.P.......K..N..R........T.........V.........E...   193
LBV         QT.R....AG.SR.S..QS.......L.....L..T.V........E...   193
LLEBV       KY.AK.I.ERFRTTQT.SLV....I.I.....L..IL.............   192
SHIBV       QT.K.....C.GRIS..QS.......L.....L..TM.........E...   193
MOKV        QT.K.....G.NR.S..QS.......L.....L.............E...   193
VSV_NJ      DY.FK-------PKNKEIDSMALV.PLCQKF.DL.K.T.IL..VSLG.T.   180
```

| | | |
|---|---|---|
| **Consensus** | TILALWKDXXSXDXKX-DXVKFRDQIWGSLXVTKDFVYSQSXNCLFDRNY | |
| RABV-Tha | ........LT.V.IGK-.L...K.....V.I..........SS....... | 243 |
| ARAV | .......ELN.V.VGK-.Q...........V..........A.S...... | 242 |
| ABLV | ......R.LN.V.I.K-.Q.....S.....I...E.....NS........ | 243 |
| BBLV | ........LS.V.I.K-.Q...........V.....I....A........ | 242 |
| WCBV | ..IS..R.IIHYET.E-.RLTIK.......L...E....S.TSA...K.. | 242 |
| DUVV | ...S..R.IS.I.I.N-.L...K.....T.V.....I....A........ | 242 |
| TWBLV | ...S..R.IA.I.TRV-.L..........VV..........S........ | 242 |
| GBLV | ......RELN.I.T.K-.Q...........I..........S.S...... | 242 |
| EBLV1 | .......EIA.I.V.S-.L...........VI.........A........ | 242 |
| EBLV2 | ........LN.V.I.K-.Q...........L..........A........ | 242 |
| IKOV | A.I...RSFLDYNSRT-NS.SVK.LL..RMV...EY.LMLDIS......F | 241 |
| IRKV | ........IA.I.V.N-.L...........V..........A........ | 242 |
| KHUV | ........LN.V.I.K-.Q...........V..........A........ | 242 |
| KBLV | ......R.LN.V.T.K-.Q...........L..........A........ | 242 |
| LBV | ........IT.V.I.N-.K.L.K.PL..KFL........YDS.S...... | 242 |
| LLEBV | ..I...RSLLDYNS.T-.S.SVK.IL..QMI....YLFIIDV.S.....F | 241 |
| SHIBV | ..I....EIT.I.V.N-.K.T...PL..KFL...E....YTNCS...... | 242 |
| MOKV | ......R.IT.I.V.N-.R.S.K.PL..K.L.......AHNN.....K.. | 242 |
| VSV_NJ | EL.TTF.GKYRMSCENIPIARL.LPSL.PVFMC.GWT.IHKERV.M...F | 230 |

| | | |
|---|---|---|
| **Consensus** | TLMLKDLFLSRFNSLLILLSPPEPRYSDDLXSXLCQLYIAGDXVLSXCGN | |
| RABV-Tha | ...............M..............I.Q.........Q...M... | 293 |
| ARAV | ..............................V.Q.........NI..I... | 292 |
| ABLV | ..............................I.L.........H...M... | 293 |
| BBLV | ..............................I.Q.........H..AM... | 292 |
| WCBV | V..............M..I....SK...T.IET..S..V...Q.VAQ... | 292 |
| DUVV | ..............F........A......V.N......S..R...V... | 292 |
| TWBLV | ..............F........SK.....V.S.........K..AS... | 292 |
| GBLV | ..............................V.QM........N...M... | 292 |
| EBLV1 | ..................I....S......V.N.........K...A... | 292 |
| EBLV2 | ...........................E..I.Q.........N...T... | 292 |
| IKOV | M.....T........S......DSL..A.FSES..D..NV..NIIAE... | 291 |
| IRKV | .......................F......V.T.........K..AS... | 292 |
| KHUV | ..............................I.Q.........N...M... | 292 |
| KBLV | ..............................I.Q.........N...T... | 292 |
| LBV | ..................I.........E.VAN..R......KI..T... | 292 |
| LLEBV | V.....T........M......DTL..S.FPENI.S..M...S..AD... | 291 |
| SHIBV | ..................I...........I.N.....KS..K.I.E... | 292 |
| MOKV | ..................V...DS......AAN..R...S..RL..T... | 292 |
| VSV_NJ | L..C..VIIG.MQTF.SMIGRSDNKF.P.QIYT.ANV.RI..KI.EQ... | 280 |

```
Consensus  SGYDVIKMLEPYVVNXLVQRAEXFRPLIHSLGDFPXFIKDKXXQLEGTFG
RABV-Tha   ...E...I.......S......K............V.....VS.......  343
ARAV       ...............S...K..E............I.....VG.......  342
ABLV       .............I.S......E............L..RE.VG.......  343
BBLV       ...............S......G............L.....VT....I..  342
WCBV       ...........C..HE..E...KY....T......E..RE.KA..I.I..  342
DUVV       ..F....L....I..K......K............Q.....TT.......  342
TWBLV      .......L.......R......E............Q..R..TH.......  342
GBLV       ...E...........S......G............V.....VG.......  342
EBLV1      .......L....I..K...K..K............Q..R..TN.......  342
EBLV2      ...............S......G...M........T.....VS.......  342
IKOV       A...I......F...K..KS..E...M.PK.....E.....TQ..I....  341
IRKV       ............I..K......S............Q..R..TT....I..  342
KHUV       ...............S......G............L.....VT.......  342
KBLV       ...............S......G............L.....VN.......  342
LBV        P...I..........L...K..T......P..E..T.....TA..I....  342
LLEBV      A..........FI..K...E..NY....PK.....E.....TR..V....  341
SHIBV      .......I.......H...K..T............A.....TT..R....  342
MOKV       A..........C..DL...R..T.........E..A..R..TI..I....  342
VSV_NJ     RA..L...I..ICNLKMMEL.RAH..K.PKFPH.EEHV.GSVRE.TQRSN  330
```

```
Consensus  PSAXXFFXXLDQLDNIHDLVFVYGCYRHWGHPYIDYRKGLSKLYDQVHXK
RABV-Tha   ...KR..RV...F...................................I.  393
ARAV       ...RK..RV.......................................V.  392
ABLV       ...RR..QV...V...........................L.......V.  393
BBLV       ...RN..HI.......................................L.  392
WCBV       .P.SR..SVI.R....................................M.  392
DUVV       ...KE..QTM.S...V.....................R..........L.  392
TWBLV      ...KE..QS..L....................................V.  392
GBLV       ...RK..QV...F...................................I.  392
EBLV1      ...RE..QTM.L....................................V.  392
EBLV2      ...RN..FV.......................................V.  392
IKOV       .V.NF..SK.E.YN..................H...............M.  391
IRKV       .R.RE..QTM.L.........T..........................V.  392
KHUV       ...RN..RI.........................E.............V.  392
KBLV       ...RN..RV.......................................V.  392
LBV        .C.SQ..SM.S.F........................R..T..F....MR  392
LLEBV      .V.DS..SQ..RFN..................H...............M.  391
SHIBV      .C.SQ..SA...F...........................T..F....M.  392
MOKV       .C.SQ..FM.Q.F........I..................T..F....M.  392
VSV_NJ     -RIQTLYDLIMSMKDVDLVLV...SF......F...FE..E..HT..NME  379
```

| | | |
|---|---|---|
| **Consensus** | **KXIDXXYQECLASDLAKRILRWGFDKYSKWYLDXXLLXXDHPLXPYIKTQ** | |
| RABV-Tha | .V..KS..........R..............I.SRF.PR....T...... | 443 |
| ARAV | .V..GD...........................AK..TK....T...... | 442 |
| ABLV | .M..GA...........................PK..AP....A...... | 443 |
| BBLV | .V..KD...........................SK..AH....T..V... | 442 |
| WCBV | .F..EG..KS.......................PARMRR....RQ.VS.. | 442 |
| DUVV | .T..KS......................R..V.PN..QN....V..VR.. | 442 |
| TWBLV | .V..QS............V.........R....TH..QN....T..V... | 442 |
| GBLV | .V..GT...........................SK..SK....A...... | 442 |
| EBLV1 | .I..RN......................R....SN..PG....S..V... | 442 |
| EBLV2 | .V..GD...........................PK..EK....I...Q.. | 442 |
| IKOV | .S..SR...S.......K.............V.VKKVPLH...RS..L.. | 441 |
| IRKV | .V..QH......................R....QE..PK....T..V... | 442 |
| KHUV | .V..GD...........................SK..APN...T...... | 442 |
| KBLV | .V..GD...........................SK.MPQ....T...... | 442 |
| LBV | .T..QK...........K...............TNS.PKN...A...A.. | 442 |
| LLEBV | .S..TS...S......R.V.........R..V.SRG.PSH...K...S.. | 441 |
| SHIBV | .T..KH...........K..........R....TS..PKN...I...T.. | 442 |
| MOKV | .T..QQ...R......RK...............TGVIPK....A...A.. | 442 |
| VSV_NJ | .H..KE.PQQ......RLV.NKQ.SESK..FV.PSKMSPK..FYEHVINK | 429 |

| | | |
|---|---|---|
| **Consensus** | **TWPPKHIVDXVGDTWHXLPITQIFEIPESMDPSEILDDKSHSFTRTKLAS** | |
| RABV-Tha | .........L......K.............................R... | 493 |
| ARAV | ......V..I..N...K................................. | 492 |
| ABLV | .........L..N...K................................. | 493 |
| BBLV | .........I..N...K................................. | 492 |
| WCBV | .........L......E.........................L..S...G | 492 |
| DUVV | ......V..V......N................................. | 492 |
| TWBLV | ....R.V..V..N...N.............................R... | 492 |
| GBLV | ......V..L..N...K.....L..V........................ | 492 |
| EBLV1 | ......V..M......S.............................R... | 492 |
| EBLV2 | .........I..N...K................................. | 492 |
| IKOV | .......A.MI.N...E.....L.D....................S..IA | 491 |
| IRKV | ......V..LI..S..N.....L........................... | 492 |
| KHUV | .........M..N...K................................. | 492 |
| KBLV | .........I..N...R................................. | 492 |
| LBV | .........LL..S..L..M.........................S...Y | 492 |
| LLEBV | .........MM.....E.....L...................L..NR.I. | 491 |
| SHIBV | ....R.V..LL..S..S..M..........................R... | 492 |
| MOKV | ......V..LL..S..T..M.....V......................S. | 492 |
| VSV_NJ | ...TAAKIQDF..N..K..LI.C....DLI...V.YS.....MNKKEVIQ | 479 |

| | | |
|---|---|---|
| **Consensus** | WLSENRGGPVPSEKVIITALSRPPVNPREFLKSIDXGGLPDDDLIIGLKP | |
| RABV-Tha | .....................K.............L.....E........ | 543 |
| ARAV | ...............................R...I.............. | 542 |
| ABLV | ..A..................K.............L.............. | 543 |
| BBLV | ...D...............................L.............. | 542 |
| WCBV | ..A..................KS........R.V.AH..DEE........ | 542 |
| DUVV | ....H..........................R...VS............. | 542 |
| TWBLV | ...............................RA..I.............. | 542 |
| GBLV | .........I.........................L.............. | 542 |
| EBLV1 | ...............................R...V.............. | 542 |
| EBLV2 | ...DH..............................L.............. | 542 |
| IKOV | ....G................QN.........R..DH..DP......... | 541 |
| IRKV | ...............................R...V.............. | 542 |
| KHUV | ...D...............................I.............. | 542 |
| KBLV | ...................................L.............. | 542 |
| LBV | ...............................RT..Q....E......... | 542 |
| LLEBV | ....GK...............QN.I.......R..DH..DQ......... | 541 |
| SHIBV | .....................KA.........AV.LN..AE......... | 542 |
| MOKV | ....H..............................Q.............. | 542 |
| VSV_NJ | HVRSKPNV.I..R..LQ.M.TNRAT.WKA...D..EN.MD.........G | 529 |

| | | |
|---|---|---|
| **Consensus** | KERELKIEGRFFALMSWNLRLYFVITEKLLANYILPLFDALTMTDNLNKV | |
| RABV-Tha | .................................................. | 593 |
| ARAV | .................................................. | 592 |
| ABLV | .................................................. | 593 |
| BBLV | .................................................. | 592 |
| WCBV | ................................H................. | 592 |
| DUVV | ...............................T.......S.......... | 592 |
| TWBLV | ...............................T.................. | 592 |
| GBLV | .................................................. | 592 |
| EBLV1 | ...............................T.................. | 592 |
| EBLV2 | .................................................. | 592 |
| IKOV | ................................H.I....S.......... | 591 |
| IRKV | ......V........................T.................. | 592 |
| KHUV | .................................................. | 592 |
| KBLV | .................................................. | 592 |
| LBV | ................................H................. | 592 |
| LLEBV | ...............................SH.I....S.......... | 591 |
| SHIBV | ................................H................. | 592 |
| MOKV | .......D........................H.I............... | 592 |
| VSV_NJ | .......A....S....R..E......Y.IKT.YV...KG...A.D.TS. | 579 |

| | | |
|---|---|---|
| **Consensus** | FKKLIDRVTGQGLXDYSRVTYAFHLDYEKWNNHQRLESTKDVFSVLDXVF | |
| RABV-Tha | .............L.........................E.......Q.. | 643 |
| ARAV | .............L.................................Y.. | 642 |
| ABLV | .............Q.............................P...Q.. | 643 |
| BBLV | .............Q.................................H.. | 642 |
| WCBV | .............M.....................M...........K.. | 642 |
| DUVV | .............L.........................I.......K.. | 642 |
| TWBLV | .............L.................................K.. | 642 |
| GBLV | .............L.................................QA. | 642 |
| EBLV1 | .............L.................................K.. | 642 |
| EBLV2 | .............Q.................................Y.. | 642 |
| IKOV | .............T.....................M...EH..Q...K.. | 641 |
| IRKV | .............L.................................KA. | 642 |
| KHUV | .............Q.................................Y.. | 642 |
| KBLV | .............Q.................................Y.. | 642 |
| LBV | .............L.................................KA. | 642 |
| LLEBV | .............S.....................M...RN..E...K.. | 641 |
| SHIBV | .............C.................................KA. | 642 |
| MOKV | .............K.................................RA. | 642 |
| VSV_NJ | I..MM.SSS....D...S.CL.N.I..........K..NGPI.R.MGQFL | 629 |

| | | |
|---|---|---|
| **Consensus** | GLKXVFSRTHEFFQKSWIYYSDRSDLIGLWXDQIYCLDMSXGPTCWNGQD | |
| RABV-Tha | ...R..........................E.........N......... | 693 |
| ARAV | ...R.............V............E........TN......... | 692 |
| ABLV | ...K..........................E.........D......... | 693 |
| BBLV | ...R.............V............EG........D......... | 692 |
| WCBV | ..ER.................E......IAGGN......TG......... | 692 |
| DUVV | ...K..........................R.........N......... | 692 |
| TWBLV | ...K..........................K........GD......... | 692 |
| GBLV | ...K..........................E.........N......... | 692 |
| EBLV1 | ...K.............V............R........TN......... | 692 |
| EBLV2 | ...K.............V............E.........D......... | 692 |
| IKOV | ...K.................E......I.GNK.C.....E.....D... | 691 |
| IRKV | ...R........................M.K.........N......... | 692 |
| KHUV | ...K.............V............E.........N......... | 692 |
| KBLV | ...R.............V............E.........N......... | 692 |
| LBV | ..SR.............V..........I.K........AE......... | 692 |
| LLEBV | ...K.................E......V.G.K.C.....E.....D... | 691 |
| SHIBV | ..SR.............V..........I.N........AD......... | 692 |
| MOKV | .MRR........................I.K........TE......... | 692 |
| VSV_NJ | .YPSLIE......E..L...NG.P..MTVRNGTLCN--STKHRV.....K | 677 |

| | | |
|---|---|---|
| **Consensus** | GGLEGLRQKGWSLVSLLMIDRESQTRNTRTKILAQGDNQVLCPTYMLSSG | |
| RABV-Tha | ................................................P. | 743 |
| ARAV | .................................................. | 742 |
| ABLV | ...............................DF................. | 743 |
| BBLV | .................................................. | 742 |
| WCBV | ............MI..........I.....................V... | 742 |
| DUVV | ................................................P. | 742 |
| TWBLV | .................................................. | 742 |
| GBLV | .................................................. | 742 |
| EBLV1 | .................................................. | 742 |
| EBLV2 | .................................................. | 742 |
| IKOV | ........................I....................VV.Q. | 741 |
| IRKV | .................................................. | 742 |
| KHUV | .................................................. | 742 |
| KBLV | .................................................. | 742 |
| LBV | ...................E.............................. | 742 |
| LLEBV | ........................I...K................VV.P. | 741 |
| SHIBV | .................................................. | 742 |
| MOKV | ...................E...K.......................... | 742 |
| VSV_NJ | ............I.N..V.Q..AKI...AV.V........I.TQ.KTKKT | 727 |

| | | |
|---|---|---|
| **Consensus** | LXXEGLLYELESISRNAXSIYRAIEDGASKLGLIIKKEETMCSYDFLIYG | |
| RABV-Tha | .SR..............L.......E........................ | 793 |
| ARAV | .TQ.......D......L................................ | 792 |
| ABLV | .SH..............L................................ | 793 |
| BBLV | .TE..............L................................ | 792 |
| WCBV | .SQD..K....N..K..M.......E..GR.................... | 792 |
| DUVV | .NK.......D......I..........A..................... | 792 |
| TWBLV | .NN.......D...K..I................................ | 792 |
| GBLV | .SH..............L................................ | 792 |
| EBLV1 | .NK.......D......I................................ | 792 |
| EBLV2 | .TQ...I...D......L.......E........................ | 792 |
| IKOV | .NED..K....N.....L.......E..R..................... | 791 |
| IRKV | .TK.......D......I................................ | 792 |
| KHUV | .TQ.......D......L................................ | 792 |
| KBLV | .TQ.......D......L................................ | 792 |
| LBV | .NN...R.......K..M.......E.................F...... | 792 |
| LLEBV | .NED..R....N.....I.......E..G..............F...... | 791 |
| SHIBV | .NN...M.......K..M.......E........................ | 792 |
| MOKV | .NN...M....N..K..M.......E.................F...... | 792 |
| VSV_NJ | RSELE.RAV.HQMAG.NNK.MEE.KR.TE......NDD...Q.A.Y.N.. | 777 |

```
Consensus  KTPLFRGNILVPESKRWARVSCISNDQIVNLANIMSTVSTNALTVAQHSQ
RABV-Tha   ..................................................  843
ARAV       ..................................................  842
ABLV       ..................................................  843
BBLV       ..................................................  842
WCBV       ..................................................  842
DUVV       ..................................................  842
TWBLV      ..................................................  842
GBLV       ..................................................  842
EBLV1      ..................................................  842
EBLV2      ..................................................  842
IKOV       ....Y.......................IS....................  841
IRKV       ..................................................  842
KHUV       ..................................................  842
KBLV       ..................................................  842
LBV        ..................................................  842
LLEBV      ............................IS....................  841
SHIBV      ..................................................  842
MOKV       ..................................................  842
VSV_NJ     .I.I...V.RGL.T...S..T.VT....PTC..L..S.........HFAE  827
```

```
Consensus  SLIKPMRDFLLMSVQAVFHYLLFSPILKDRVYKILXAX-GDXFLLAMSRI
RABV-Tha   ............................G......S.E-..N........  892
ARAV       ...................................S.D-..E........  891
ABLV       .......G....A...............G......S.E-..D........  892
BBLV       ...................................S.D-..D....L...  891
WCBV       ................IY.................N.Q-.ED..MS.A..  891
DUVV       ............................N......V.D-.TE........  891
TWBLV      ...................................A.E-.TE........  891
GBLV       ............................G......S.E-..D........  891
EBLV1      .................Y.................V.D-.TE........  891
EBLV2      ............A......................S.E-..N........  891
IKOV       ..V..I....M......Y..F.......S..HC..NLS-HERL.......  890
IRKV       ..V................................V.D-.NE........  891
KHUV       ...................................S.D-..D........  891
KBLV       ...................................S.D-..D........  891
LBV        ................IY...............V.NSK-..D........  891
LLEBV      ..V..I..........IY..........S..HG..SLS-.ERL....A..  890
SHIBV      ................IY...............V.NSK-..D...T....  891
MOKV       ..V.............IY........I......V.NSK-D.D........  891
VSV_NJ     NP.NA.IQYNYFGTF.RLLLFMHD.AIRQSL.NVQEKIP.LHTRTFKYAM  877
```

| | | |
|---|---|---|
| **Consensus** | IYLDPSLGGVSGMSLGRFHIRQFSDPVSEGLSFWKEIWXSSSEXWIHSLC | |
| RABV-Tha | ..................................R...L..H.S...A.. | 942 |
| ARAV | ......................................S....P..Q... | 941 |
| ABLV | V.....................................T....S...... | 942 |
| BBLV | V..............................A......S....A...... | 941 |
| WCBV | ...........................T..........EV...G...... | 941 |
| DUVV | ..................................R...S....T...... | 941 |
| TWBLV | ......................................L..N.P...... | 941 |
| GBLV | ......................................A....S...A.. | 941 |
| EBLV1 | ..................................R...L....S...A.. | 941 |
| EBLV2 | ...............................A......S....S...... | 941 |
| IKOV | .F.........................T..........EC...E..L... | 940 |
| IRKV | V..............................A..R...L..N.P...... | 941 |
| KHUV | VF....................................S....S...... | 941 |
| KBLV | ......................................S....A...... | 941 |
| LBV | ...............................T......L.A..T...H.. | 941 |
| LLEBV | .........I.................T..........E....N..LA.. | 940 |
| SHIBV | ...............................T......L....S...H.. | 941 |
| MOKV | ...............................T......L....A.V.H.. | 941 |
| VSV_NJ | L.....I...C..A.S..L..A.P...T.S.....F.YEHA..PHLKKMA | 927 |

| | | |
|---|---|---|
| **Consensus** | QEAGNPDLGDRSLESFTRLLEDPTTLNIRGGASPTILLKEAIRKALYDEV | |
| RABV-Tha | .........E.T................K..........D.......... | 992 |
| ARAV | .................................................. | 991 |
| ABLV | ..............K........................D.......... | 992 |
| BBLV | .......................................D.......... | 991 |
| WCBV | ...................I.............................. | 991 |
| DUVV | .........E........................................ | 991 |
| TWBLV | ...........T...................................... | 991 |
| GBLV | .........E..................K..........D.......... | 991 |
| EBLV1 | ......................................R........... | 991 |
| EBLV2 | .................................................. | 991 |
| IKOV | .......I.E.T...............................Q...... | 990 |
| IRKV | ...............................................E.. | 991 |
| KHUV | .................................................. | 991 |
| KBLV | .................................................. | 991 |
| LBV | ...........T...................................... | 991 |
| LLEBV | .......I.E.T...............................Q...... | 990 |
| SHIBV | .................................................. | 991 |
| MOKV | .................................................. | 991 |
| VSV_NJ | VMF.D.PIAKFRI.HINK......S...SM.M..AN...SEVK.C.IESR | 977 |

| | | |
|---|---|---|
| **Consensus** | **DKVENSEFREAILLSKTHRDNFILFLKSIEPLFPRFLSELFSSSFLGIPE** | |
| RABV-Tha | ............................V..................... | 1042 |
| ARAV | ............................V..................... | 1041 |
| ABLV | ...............RH................................. | 1042 |
| BBLV | ..........................R.V..................... | 1041 |
| WCBV | .R................................................ | 1041 |
| DUVV | ..........................R.V..................... | 1041 |
| TWBLV | ............................V..................... | 1041 |
| GBLV | .................................................. | 1041 |
| EBLV1 | ..........................R.V..................... | 1041 |
| EBLV2 | .................................................. | 1041 |
| IKOV | ...S........I.........V.Y.R....................... | 1040 |
| IRKV | ............................V..................... | 1041 |
| KHUV | ..........................R....................... | 1041 |
| KBLV | ..........................R....................... | 1041 |
| LBV | ............I..................................... | 1041 |
| LLEBV | ...S........I.............R....................... | 1040 |
| SHIBV | ............I..................................... | 1041 |
| MOKV | .R..........I..................................... | 1041 |
| VSV_NJ | SSIK.EIIKD.TIYMHQEEEKLRG..W..K.........FKAGT...VS. | 1027 |

| | | |
|---|---|---|
| **Consensus** | **SIIGLIQNSRTIRRQFRRSLSRTLEESFFNSEIHGINRMTQVPQRIG--R** | |
| RABV-Tha | ............................Y............T...V.--. | 1090 |
| ARAV | ................................V........I.....--. | 1089 |
| ABLV | ......................S.....Y.......S..........--. | 1090 |
| BBLV | ................................V..............--K | 1089 |
| WCBV | .....................QS......K..L..LT.L......V.--. | 1089 |
| DUVV | .................K...........S...Q..S.V...A.K..--. | 1089 |
| TWBLV | ......................A.............S.......K..--. | 1089 |
| GBLV | ............................Y..................--. | 1089 |
| EBLV1 | ....................................S..........--. | 1089 |
| EBLV2 | .............................................V.--. | 1089 |
| IKOV | .................K.......K..YK..LS.LEK.A.MG..V.--T | 1088 |
| IRKV | ......................A............V...........--. | 1089 |
| KHUV | .................K..............V..............--. | 1089 |
| KBLV | ...............................................--. | 1089 |
| LBV | ...........V.....K....A......H...........T...L.--. | 1089 |
| LLEBV | .................K..........YR..LN.LE....TG..V.--. | 1088 |
| SHIBV | ...........V.....K....A......H......S....S...L.--. | 1089 |
| MOKV | ...........V.....K..........LQ........V..T...L.--. | 1089 |
| VSV_NJ | GL.N.F.......NS.KKRYHKD.D.LIIK...SSLSHLGSMHY.L.DNQ | 1077 |

```
Consensus   VWXCSAERADXLREISWGRKVVGTTVPHPSEMLXLXPKSSISCXCGXXGX
RABV-Tha    ..P..S....L......................G.L.R.....T..TT.G   1140
ARAV        ..D.......Q..................A...A.L.......T..QT.E   1139
ABLV        ..S..S...HL.........R............A.L.......P..LT.S   1140
BBLV        ..A.......L..................A...V.V.R.....V.TQT.E   1139
WCBV        ..R.......Q.......K..............N.V....V..N..RINP   1139
DUVV        ..R.......Q......................SMI.......T..QLED   1139
TWBLV       ..N.......Q......................VMV.......N..QS.E   1139
GBLV        ..S..S....L......................SML.......T..SS.G   1139
EBLV1       ..S.......Q......................A.I.......T..QS.D   1139
EBLV2       ..N.......L..................G...V.L....V..V.KQT.E   1139
IKOV        ..K.......R...................L..S.T.R.....G.RIQDV   1138
IRKV        ..A.......Q.....................IA.I.......T.SQS.E   1139
KHUV        ..P.......L......................A.L.......V..QT.E   1139
KBLV        ..T.......L..................A...V.L.......I..QT.E   1139
LBV         ..P.......Q..................A...V.V.R..VA.G.NVRDA   1139
LLEBV       ..D..SDK..R......................S.T....V..G.KISDI   1138
SHIBV       ..S.......Q......................M.V....V..G.SIREL   1139
MOKV        ..A.......Q..........I...........A.I....VA.G.YTREA   1139
VSV_NJ      I.S...S...V..YK..T...........L..HGPPS.KEKP.HLCNS-S   1126
```

```
Consensus   XXPRISVSVLPSFDQSFFSRGPLKGYLGSSTSMSTQLFHAWEKVTNVHVV
RABV-Tha    EN..V.............................................   1190
ARAV        NS................................................   1189
ABLV        EN..V.I.....IY......................T.............   1190
BBLV        DS............R...................................   1189
WCBV        ES............N..L..............VT............I...   1189
DUVV        DN.............T..................................   1189
TWBLV       DN............H..L................................   1189
GBLV        EN..V...........................T.................   1189
EBLV1       DS..........L.....................................   1189
EBLV2       DS................................................   1189
IKOV        QK..V.........NR.V..............V.................   1188
IRKV        AN..............Y.................................   1189
KHUV        DS................................................   1189
KBLV        DS................................................   1189
LBV         YN..L.........N.................V.................   1189
LLEBV       IK..V.........CR.D..............V.................   1188
SHIBV       YN............N......S.R........V.................   1189
MOKV        GN............P..L.....................S..........   1189
VSV_NJ      GLTY...HCPKGITDV.NR....PA....N..E..SILQP...ESKIPII   1176
```

| | | |
|---|---|---|
| **Consensus** | **KRALSLKESINWFIXRDSNLAQTLIRNIXSLTGPXFPLEEXPVFKRTGSA** | |
| RABV-Tha | ..............T.N...........M.....D.....A......... | 1240 |
| ARAV | ..............V..........K..I.....S.....T......... | 1239 |
| ABLV | ..............S.N...........I.....Q.....A......... | 1240 |
| BBLV | ..............D.K........K..I.....L.....T......... | 1239 |
| WCBV | R.............D......KI.....L.....D.VI..T.L....... | 1239 |
| DUVV | ..............S......H......V.....S.....A......... | 1239 |
| TWBLV | ..............I..........K..M.....A..I..A......... | 1239 |
| GBLV | ..............V.............M.....D.....A......... | 1239 |
| EBLV1 | ..............T.............V.....F.....A......... | 1239 |
| EBLV2 | ..............A.............T.....Q.....T......... | 1239 |
| IKOV | R.............E......KI.....K.....E.EID.I.L....... | 1238 |
| IRKV | ..............T..........K..V.....T.....A......... | 1239 |
| KHUV | ..............V.......V..K..I.....L.....T......... | 1239 |
| KBLV | ..............V..........K..I.....Q.....T......... | 1239 |
| LBV | .............VS.............L.....S..I..A......... | 1239 |
| LLEBV | R.............N......K......K.....E.EI..I......... | 1238 |
| SHIBV | ..............S.............L.....E..I..A......... | 1239 |
| MOKV | .............VS......K...G..L.....S..I..T......... | 1239 |
| VSV_NJ | ...TR.RDA.S...PPE.P.STCILN..RA...EDWSSK-QHG....... | 1225 |

| | | |
|---|---|---|
| **Consensus** | **LHRFKSARYSEGGYSSICPNLLSHISVSTDTMSDLTQDGXNFDFMFQPLM** | |
| RABV-Tha | ................V......................K.Y........ | 1290 |
| ARAV | .......................................T.......... | 1289 |
| ABLV | ....................................H..R.......... | 1290 |
| BBLV | .......................................R.......... | 1289 |
| WCBV | ................V......................S.......... | 1289 |
| DUVV | .......................................T.......... | 1289 |
| TWBLV | ....................M..................T.......... | 1289 |
| GBLV | ................V......................R.......... | 1289 |
| EBLV1 | .......................................T.......... | 1289 |
| EBLV2 | .......................................K.......... | 1289 |
| IKOV | ......S.........V................E..H..R.......... | 1288 |
| IRKV | .......................................T.......... | 1289 |
| KHUV | .................................E.....K.......... | 1289 |
| KBLV | .......................................K.......... | 1289 |
| LBV | ................V......................T.......... | 1289 |
| LLEBV | ................V................E..HG.L.......... | 1288 |
| SHIBV | ...............AV......................T.......... | 1289 |
| MOKV | ...............AV......................T.......... | 1289 |
| VSV_NJ | ....STS.M.N..FA.QS.AT.TRMIAT....R.FG--TK.Y.....ASL | 1273 |

```
Consensus  LYAQTWTSELVQKDLRLXDSTFHWHLRCXKCIRPIDDIXLEAPQVFXFPD
RABV-Tha   ............R.T..K..........NR.V......I..TS.I.E...  1340
ARAV       ..............D..K..........P.........N.......M...  1339
ABLV       ..............I..K..........P....S...VI.VH....D...  1340
BBLV       ..............I..R..........L.......E.T.....I.T...  1339
WCBV       ..S......I..R....R.T.Y......QR.....E..T.DS.CC.V...  1339
DUVV       .................K..........VR.V......T....N..S...  1339
TWBLV      .................K..........LR.V.....VV....N..S...  1339
GBLV       ..............T..R..........H..V......T..T....A...  1339
EBLV1      .................R..........L.........T....KI.S...  1339
EBLV2      ..............I..R..........L.........I.D.....M...  1339
IKOV       ..S....A....RN.K.R...Y......KD.V.....VV..C.KI.E...  1338
IRKV       .................K..........L........VI..T....V...  1339
KHUV       .................K..........P.........T.......M...  1339
KBLV       ..............I..R..........Q..V......T.......M...  1339
LBV        ..............V..K..........Q......EEVS.....S.I...  1339
LLEBV      ..S....A.I..R..K.R...Y......HN...I...VT..C....D...  1338
SHIBV      .................R..........Q......EEVS.....S.A...  1339
MOKV       .................A..........Q......EEVT.D...L.A...  1339
VSV_NJ     ..G.MT..IS--RYGNPGSC.D.Y.I..KG...E.EEVE.NTSLEYKT..  1321
```

```
Consensus  VSKRISRMVSGAVPQFXKLPEIXLKPGXFEXLXGKXKSRHIGTAQGLLYS
RABV-Tha   ..............H.Q...D.R....D..S.S.RE......S.......  1390
ARAV       ..............H.Q.....N....K..L.S..D..............  1389
ABLV       ..............SSQI...VN.L..H..S.C.RD......S.......  1390
BBLV       ................QR...VV....K..S.N..E..............  1389
WCBV       I.N...........EVR....VA..A.D.STISPRER.Y...........  1389
DUVV       ..R.............Q.....S....K..T.S..DQ.K...........  1389
TWBLV      ................R.....E....K.DT.S..D..............  1389
GBLV       ................Q.....H.RA.H..S.G..E......S.......  1389
EBLV1      ................R.....G....K.DS.KE.D..............  1389
EBLV2      ................QR....N....K..A.DS.D..............  1389
IKOV       I.T...........E.K....VS..Q.NLSMMSNED..F...........  1388
IRKV       ................R...D.T....K.DA.K..E..Q...........  1389
KHUV       ................QR....S....K..S.N..D..............  1389
KBLV       ................Q.....N....K..P.S..D..............  1389
LBV        I.T.............R...D.G..A.DLTV.NNNER.Y...........  1389
LLEBV      I.T...........E.K.....H.Q..DLNTMTD.S..F...........  1388
SHIBV      I.T.............R...T.E..A.DLSS.TNNER.Y.V.........  1389
MOKV       I.S.............R....VG..A.DLMA.SSSER.Y...........  1389
VSV_NJ     .YHILEKWRNNTGSWGHQIKQLKPAE.NW.S.SPVEQ.YQVARCI.F..G  1371
```

| | | |
|---|---|---|
| **Consensus** | ILVAXHDSGYNDGTIFPVNIYXKVSPRDYLRGLARGVLIGSSICFLTRMT | |
| RABV-Tha | ....I.......S........S............................ | 1440 |
| ARAV | V...T................GR........................... | 1439 |
| ABLV | ....T..P.............S.I..K.........I.......L..... | 1440 |
| BBLV | ....T..A.............S............................ | 1439 |
| WCBV | ....I..P....HAL......SRM..QH.....S................ | 1439 |
| DUVV | ....I.........V......KR........................... | 1439 |
| TWBLV | ....T.........L......KR....E...................... | 1439 |
| GBLV | ....T................G............................ | 1439 |
| EBLV1 | ....T................G............................ | 1439 |
| EBLV2 | ....I.......A....M...S.I.........S..I............. | 1439 |
| IKOV | ....I..P....HS.......SRL...G........I............. | 1438 |
| IRKV | ....T................G............................ | 1439 |
| KHUV | ....T................G....K....................... | 1439 |
| KBLV | ....T................G............................ | 1439 |
| LBV | ....V..P....NSL......GR..A.S...................... | 1439 |
| LLEBV | ....I..P....H.L......P.L.AKE..L..S..I............. | 1438 |
| SHIBV | ....I..P....NSL......G...A.S.........F............ | 1439 |
| MOKV | ....V..P....NSL......G...A.G........I............. | 1439 |
| VSV_NJ | E.THKKSRQAD.SSL..LS.QL..RG.GF....LD.LMRS.CCQVIH.RS | 1421 |

| | | |
|---|---|---|
| **Consensus** | NININRPLELISGVISYILLRLDNHPSLYVMLKEPSLRSEIFSIPQKIPA | |
| RABV-Tha | .............................I..R.....G........... | 1490 |
| ARAV | .................................................. | 1489 |
| ABLV | ...............T................R................. | 1490 |
| BBLV | ..................................S............... | 1489 |
| WCBV | ....................K..T...........GM............. | 1489 |
| DUVV | S.....................................L........... | 1489 |
| TWBLV | .................................................. | 1489 |
| GBLV | ................................R................. | 1489 |
| EBLV1 | .................................................. | 1489 |
| EBLV2 | ................................R................. | 1489 |
| IKOV | K..V......M............S........R..LI..........V.. | 1488 |
| IRKV | ...............................................V.. | 1489 |
| KHUV | .................................................. | 1489 |
| KBLV | ..................................Q............... | 1489 |
| LBV | ...................................D..A........... | 1489 |
| LLEBV | ...V......M.........K..S.....M....SDI............. | 1488 |
| SHIBV | ...................................E.............. | 1489 |
| MOKV | ....................K........I.....E..A........V.. | 1489 |
| VSV_NJ | VSTLK..ANAVY.GLI.LIDK.SASSPFLSLVRTGPI.Q.LEQV.H.MST | 1471 |

| **Consensus** | **AYPTTMKEGNRSVLCYLQHVLRYERXXITXSPENDWLWIFSDFRSXKMTY** | |
|---|---|---|
| RABV-Tha | ............I............EV..A...............S.... | 1540 |
| ARAV | .........................DA..A..D............T.... | 1539 |
| ABLV | .........................EI..S...............M.... | 1540 |
| BBLV | .........................DM..S..D............A.... | 1539 |
| WCBV | ...........AI.S...YI...QKEE..R.ESS.L..M......M.... | 1539 |
| DUVV | .........................EA..A...C...........I.... | 1539 |
| TWBLV | .......D..........Q......EA..A...G...........I.... | 1539 |
| GBLV | .........................EV..A..............NI.... | 1539 |
| EBLV1 | .........................EA..T...............I.... | 1539 |
| EBLV2 | .........................DV..A...............A.... | 1539 |
| IKOV | ......R....A..SH..YT...DKDSLVDGSGSN...V......S.... | 1538 |
| IRKV | .........................EA..A...............I.... | 1539 |
| KHUV | .........................DV..A...............T.... | 1539 |
| KBLV | .........................DV..A...............T.... | 1539 |
| LBV | ......E...........Q......DSMCS...SEL.........I.... | 1539 |
| LLEBV | ...........AI.S....T....KENLIEGSGKN...V......S.... | 1538 |
| SHIBV | ..................Q......DSMSS..G..L.........I.... | 1539 |
| MOKV | ......E...........Q......DSMSFP.G..I.........I.... | 1539 |
| VSV_NJ | S...NIRDLGSI.RN.FKYQC.PVERG-HYKTYYNQI.L...VL.TEFI- | 1519 |

| **Consensus** | **LTLITYQSHILLQKIEKNLSKQMRXXLRQLXSLMRQVLGGHGEXXLDSDE** | |
|---|---|---|
| RABV-Tha | .........L...RV.R....S..TN...MS............DT.E..D | 1590 |
| ARAV | ...V....................IR....S....R.......ET...E. | 1589 |
| ABLV | ..........F..R..RS...K..AD....S............DT...G. | 1590 |
| BBLV | ...V........L....S......IR....S............DS.E... | 1589 |
| WCBV | ...L.F.T.......G.S......SD..H.NT...R....Q..GD...Q. | 1589 |
| DUVV | .........L.....DR.......VQ....N............GS...AD | 1589 |
| TWBLV | ............L..D.G......SK....N............NI.E... | 1589 |
| GBLV | .............RV.RC...K..AS...MS............ET..... | 1589 |
| EBLV1 | .........L...R.DR.......VR....N............GT..... | 1589 |
| EBLV2 | ...V...............P....IR....S.....I......DT..... | 1589 |
| IKOV | .......T.L....VG.....K.KSD....S.........QSDMSMKDVD | 1588 |
| IRKV | .............RV.........VK....N............ST..... | 1589 |
| KHUV | ...V.............S......IR....S............DS..... | 1589 |
| KBLV | ...V..........V..S......IR....S............DT..... | 1589 |
| LBV | .......A.LW...V.R.......AK....N............DNVE... | 1589 |
| LLEBV | .....F.A....LRVS.......KAD....NT........Q..SQSKEID | 1588 |
| SHIBV | .....F.A.LW..R..R.......AK....N............ENIE..D | 1589 |
| MOKV | .....F.AYLW..RV.RS....V.IK....N............DTI...D | 1589 |
| VSV_NJ | -GPMAIS.SL.KLLYRPS.T.KD.EE..E.AA.SSNLRS.EDWDD.HIK- | 1567 |

```
Consensus  DIQXLLRDALQRTRWVDQEVRHAAKTMXXDXSPXKKXSRKAGCSEWICSA
RABV-Tha   ...R..K.S.R.............R..TGGY..N..V.........V...  1640
ARAV       ...G.......................SG.H..S.RI.............  1639
ABLV       ...R.....I...K.............TN.H..S..T...V.........  1640
BBLV       ...G.......................TG.H..SR.IP..G.....V...  1639
WCBV       .VGR..N.SVR..............SLCKEPKEPL.T..RLCQV..S..L  1639
DUVV       ..HG.....................A.KC.Y..Q.RS....SS.......  1639
TWBLV      ...G.....................N.KCEY..SR.T.............  1639
GBLV       ...R....................R..TG.Y..N.RV.........V...  1639
EBLV1      ...G.......................KC.Y..S.RV.............  1639
EBLV2      ...G.......................TG.H..S..V.............  1639
IKOV       ID.RVIC..IH.VK.I.........ELHRSEDSPSSVF.RSSQ...V..S  1638
IRKV       ...S....S.R..K.............TCEY..S..V.............  1639
KHUV       .F.G.......................TG.H..T..V.....G.......  1639
KBLV       ...G.......................TG.H..S..V.............  1639
LBV        E.HS..KE..R..............S.SP.L..IPRI...I.S.......  1639
LLEBV      VD.RI.SE.IT.VK..........R.LSPCKNSLPTLF.RP.Q......S  1638
SHIBV      E.NS..KE..R................RP.L..VP.N...I.S.......  1639
MOKV       E.LS..KES.R..............S.TP.LN.VP.I..RI.S.......  1639
VSV_NJ     -------FFSNDLLFCS..I...C.FGIKKDNEDITFYPN-WGT.Y.GNV  1609
```

```
Consensus  QQVAISTSSNPAPXSEXDVRALSRKXQNPLISGLRVVQWATGAHYKLKPI
RABV-Tha   ....V...A....V..L.I....K.F........................  1690
ARAV       .............T..L.......RF........................  1689
ABLV       ...S.........V..M...T....L............C...........  1690
BBLV       .............M..L.......RL........................  1689
WCBV       ...........SAYTDM.L.MI..QI.........I..........I..L  1689
DUVV       ..I..........T..I.I......L........................  1689
TWBLV      R...........HT..I.I......L....V...................  1689
GBLV       ........A....V..L...T....L........................  1689
EBLV1      ....V........I..I.I....K.L........................  1689
EBLV2      .............T..M.......RF........................  1689
IKOV       ..L..T..A..SHP.PI..K.I.K.L....................I...  1688
IRKV       .............V..M......K.L........................  1689
KHUV       .............T..L........F........................  1689
KBLV       .............V..L........F........................  1689
LBV        ..I.....L...LA.DI.L.S...QF....M....I..........V...  1689
LLEBV      .FIT....A..SEP.PI..KSV.KRL....................I...  1688
SHIBV      ..I.F...L....M.DI.L.L...QF....M...................  1689
MOKV       ..I.....L...SA.DI.L.S...QY....................I...  1689
VSV_NJ     IDIPVFYR---.QNVQK.-IKVPPRI....M....LG.LP......MRT.  1655
```

| **Consensus** | **LDDLXXXPXLCLVVGDGSGGISRXVLSMFPDAKLVFNSLLEVNDLMASGT** | |
|---|---|---|
| RABV-Tha | ....NVF.S..............A..N....................... | 1740 |
| ARAV | ...IDVY.A..............V..................S....... | 1739 |
| ABLV | ..N.NTY.SF.............T..N...............S....... | 1740 |
| BBLV | ....DAY.S..............T........R.........S....... | 1739 |
| WCBV | .QK.DQC.QM.............A..RV......L............A.. | 1739 |
| DUVV | .NE.DTY.T..............A..................S....... | 1739 |
| TWBLV | .NE.DSY.T..............V...L...................... | 1739 |
| GBLV | ....NSF.S..............A..N....................... | 1739 |
| EBLV1 | .N..EAY.T..............A.......................... | 1739 |
| EBLV2 | ..N.EAY.S..............T....................F..... | 1739 |
| IKOV | .NS.QFE.K..............CI.QF..GT........QL..M..... | 1738 |
| IRKV | ....DFY.T..............A.......................... | 1739 |
| KHUV | ....ESC.S..............T........R.........S....... | 1739 |
| KBLV | ....DAY.S..............T..................S....... | 1739 |
| LBV | .N..DVC.C.S..I.........V.......S.................. | 1739 |
| LLEBV | .NT.RFD.K....I.........CI.QF..GTR.......Q......... | 1738 |
| SHIBV | ....EVG.S.S............T..N....S.................. | 1739 |
| MOKV | .N..DVC.C.S..I.........V.......S.................. | 1739 |
| VSV_NJ | VSR.KISYHDF.AC......MTAAL.RHNRTSRGI.....DLS.T.LR.S | 1705 |

| **Consensus** | **HPLPPSALMSGGXDIXSRVIDFXSIWEKPSDLRNXSTWRYFQSVQXXXNM** | |
|---|---|---|
| RABV-Tha | .......I....D..I...V..E...........LT..K......KQV.. | 1790 |
| ARAV | .......IV...D..V......D...........P...K...T..CSH.. | 1789 |
| ABLV | .......I....E..T......E...........L..........SQL.. | 1790 |
| BBLV | .......I....S..V......G...........L........I.GRL.. | 1789 |
| WCBV | .........H..DELTN.....EA..........V...K......ERSK. | 1789 |
| DUVV | ........V...E..I......Q...........P..........QTT.. | 1789 |
| TWBLV | ............E..T......H...........P.......T..QAS.. | 1789 |
| GBLV | .......I....D..V......N...........L...K......NQV.. | 1789 |
| EBLV1 | ............E..V......N...........P.......T..SAA.. | 1789 |
| EBLV2 | .......IV...D..V......G...........L........I.TVN.. | 1789 |
| IKOV | ........SH..KSLTD..V..EL..........SV..N..RD..RDLST | 1788 |
| IRKV | ............E..I......D...........T..........QSI.. | 1789 |
| KHUV | .......I....D..V......E...........L..........NSSK. | 1789 |
| KBLV | .......I.R..D..V......G...........L..........SNN.. | 1789 |
| LBV | .........R..EE.T......D...........PL..K..H...SKLKT | 1789 |
| LLEBV | ........SH..P.MT......D...........I......KD..KDLGL | 1788 |
| SHIBV | .........R..E..T......E...........SV..K......TRVKA | 1789 |
| MOKV | .........R..D..T......D...........PL..K..H.I.SKLRS | 1789 |
| VSV_NJ | S.E.....ETL.GER-V.CVNGD.C..H....SDEN..K..LHLKKGCG. | 1754 |

| Consensus | SYDLIICDAEVTDIXSVNKITLLMSDFS-LSIXGPLXLIFKTYGTMLVNP | |
|---|---|---|
| RABV-Tha | ..............A.I.R........A-...D...Y.V........... | 1839 |
| ARAV | ..E...........P............A-...N...T.V........... | 1838 |
| ABLV | ..............A............V-...D..VD....S..S...D. | 1839 |
| BBLV | ..............P.............-...N...N............S | 1838 |
| WCBV | .F..V........VT.I...S......I-MA.KS.VT..........I.. | 1838 |
| DUVV | ..............S.I...........-...N...S............. | 1838 |
| TWBLV | C.............P.I...........-...N...N............. | 1838 |
| GBLV | ..............A.I.R........A-...D...Y............. | 1838 |
| EBLV1 | ..............P.I...........-...N...S............. | 1838 |
| EBLV2 | .....V........P.............-...N...N............. | 1838 |
| IKOV | NF...V........S.M.R.AF.L.N.I-...KS..T....S........ | 1837 |
| IRKV | ..............H.............-...N...N............. | 1838 |
| KHUV | ..............S.............-...N...T............S | 1838 |
| KBLV | Y.............S.............-...N...N............. | 1838 |
| LBV | QF...V........E........L....-M..R...C............. | 1838 |
| LLEBV | NF...V.......LV.....S..L...I-...KS..T............. | 1837 |
| SHIBV | HF...V........E........L...A-M..R...C..........I.. | 1838 |
| MOKV | QF...V........E........L....-M..K...C............. | 1838 |
| VSV_NJ | .IN..TM.M..Q.PAISYR.ES.VRQYVPVLLESDGC..Y.....YIATQ | 1804 |

| Consensus | DYKAIQHLSRAFPSVTGYITQMTSSFSSELYLXFSKRGKFFRDXEYLTSS | |
|---|---|---|
| RABV-Tha | E.R..............F..............R..........A...... | 1889 |
| ARAV | ................................K..........A...... | 1888 |
| ABLV | .............Q.......L..........R..........A...... | 1889 |
| BBLV | ...............A.............I..R..........S...... | 1888 |
| WCBV | ....LRN......E.......L.......I..M.K.H.R..KEA.S..A. | 1888 |
| DUVV | ..R...........AA.....V..........K..........P.H.... | 1888 |
| TWBLV | ..................T.............K..........P...... | 1888 |
| GBLV | ..........V..T...F..............R..........V...... | 1888 |
| EBLV1 | ....V...........................K..........S...... | 1888 |
| EBLV2 | ................................K..........A...... | 1888 |
| IKOV | ....LNY..QV..NMS.V...T..........LCA.S.D.YKET..I.A. | 1887 |
| IRKV | E...............................K..........P...... | 1888 |
| KHUV | ................................K..........A...... | 1888 |
| KBLV | ................................K..........A...... | 1888 |
| LBV | .....H.......N...FV..........V..K...K.H....H.L..A. | 1888 |
| LLEBV | E...LN....S...IS.V...T.......M.MIGT.S.AY.KEV..I.A. | 1887 |
| SHIBV | .....H.......NMI.FV..........V..R...K.H...EH.S..A. | 1888 |
| MOKV | .....H.......N...FV..........I..R...T.Y....H.L..A. | 1888 |
| VSV_NJ | KDNSLTLVGSL.H..QLVQ.DLS..NT.....VCKRLKDYVDT-PFVDWI | 1853 |

```
Consensus  TLREMSLVLFNCSSPKSEMXRARSLNYQDLVRGFPEEIISNPYNEMIITL
RABV-Tha   ...................Q..............................  1939
ARAV       ...................Q..............................  1938
ABLV       ..................LQ..................V...........  1939
BBLV       ...................Q..............................  1938
WCBV       .VH.........RT.....L.......S..I...................  1938
DUVV       ..............S....L..X...........................  1938
TWBLV      ..............S....L...............D..............  1938
GBLV       ...................Q..............................  1938
EBLV1      ..............S....L..............................  1938
EBLV2      ...................Q..............................  1938
IKOV       SI..L.......R......T.......K..IM...P......F.......  1937
IRKV       ..............S....L..............................  1938
KHUV       ...................Q..............................  1938
KBLV       ...................Q...............D..............  1938
LBV        .I...........NT....L...T..........................  1938
LLEBV      .I..L.......R......I.......K..TM...Q......F.......  1937
SHIBV      .I...........N.....L..........I...................  1938
MOKV       .I...........N.....L...T......I....P..............  1938
VSV_NJ     E.YDHWEKQYAFK.F.D.FN.....TPETTLI.I.PQF.PD.GVNLETLF  1903
```

```
Consensus  IDSEVESFLVHKMVDDLELQRGXLSKMSIIIAIXXVFSNRVFNVSKPLXD
RABV-Tha   ...D..................T..........MI.............T.  1989
ARAV       ......................A...I..V...MM.............N.  1988
ABLV       ......................T..R.......VI.Y...........N.  1989
BBLV       ......................T.......V..VV.............SG  1988
WCBV       ....M.......L......A..S...L.K.LV.IILY..........MR.  1988
DUVV       ..ND.........I........AF..L...LT.MV.............T.  1988
TWBLV      ..ND...............KK.AF..I...L..IM.....M.......S.  1988
GBLV       ...D..................T..........VI.............N.  1988
EBLV1      ..ND...............RQ.AF......LT.MM.............N.  1988
EBLV2      ......................T.A........VM.............T.  1988
IKOV       ...................T..S.QRLAR.L..VV.Y...........TG  1987
IRKV       ..ND..................SF......VT.ML.............N.  1988
KHUV       ....................K.A..........VM.............S.  1988
KBLV       ......................T..........VM.............S.  1988
LBV        ...............E...K..YP...A..L..TI...........S.KE  1988
LLEBV      .............I.....TK.S.Q.LAK....VI.Y......I....SS  1987
SHIBV      ............L......K..SS...A..V..IIL..........SIK.  1988
MOKV       ............I......K..SS...A.....AIL...S......S.S.  1988
VSV_NJ     QIAG.PTGVA.GITHHILQSKDK.--I.NA.GSMC.I.HFII.TI-RTT.  1950
```

```
Consensus  PXFYPPSDPKILRHFNICCSTLXYLSTALGDVXNFARLHE--LYNXPVTY
RABV-Tha   .L...................MM...M.....PS.....D--...R.I..  2037
ARAV       .K....................M.....M...LS......--...G.I..  2036
ABLV       .L...................MM....V....P....Q..--...G.I..  2037
BBLV       SV....................M.........L.......--...N....  2036
WCBV       SS.C......L...Y...S...LF.A.V...TNS.TK...--M..Q....  2036
DUVV       TK..........K.........L...V.....L.......--...N.I..  2036
TWBLV      .K....................M...V.....L....V..--...N..V.  2036
GBLV       .L...................MM.........P.......--...R.I..  2036
EBLV1      .K.................G..I...A.....L.......--...N....  2036
EBLV2      .M....................M.........L.......--...N.I..  2036
IKOV       QK.I......L.......A..FLF.NSLI..IEG.S...F--...EKIS.  2035
IRKV       .K.................G..I.........L.......--...N....  2036
KHUV       .M....................M.........L.......--...N....  2036
KBLV       .V....................M.........P.......--...N.I..  2036
LBV        SK.F......L.......S...LF...T...LP..T.I..--...L....  2036
LLEBV      HK........L.......AG.FL.I.SL...TA..T...L--...EKI..  2035
SHIBV      TK.F..............L..MLF...TM..LS..TKI..--...S..I.  2036
MOKV       .Q.F......L.......S...LF..A.M..LS..T....--...S..A.  2036
VSV_NJ     SMPG....GDVNKMCSALIG.CFW..WMES.LNLYKTCLRSIMKSM..RW  2000

Consensus  YFKKQXIXGXXYLSWSWSDXTSVFKRVACNSXLSLSSHWIRLIYKIVKTT
RABV-Tha   .....V.R.NI........D...........S..................  2087
ARAV       .....I.G.SI........NS.I........N...............R..  2086
ABLV       ...R.I.R.SI........D.......S...N..................  2087
BBLV       .....VVK.SI........S..I........N...............R..  2086
WCBV       ....RT.REKI..M....GKSP.....S..HAI...A.....L...IR..  2086
DUVV       ..R..V.G.RM..A....HS...........N....A........V....  2086
TWBLV      .....I.D.RT..G....NT.H.........C..................  2086
GBLV       .....VVR.SI........D...........N..................  2086
EBLV1      .....TLG.RM..A...T.N.P.........S..................  2086
EBLV2      .....V.R.SI......C.T.....K.....N...............R..  2086
IKOV       .....THK.KV..V....NESPTC.Q.....M.........M...VIRV.  2085
IRKV       .....ILG.RM......A.N.PI........S..................  2086
KHUV       .....V.R.SI........S...........N...............R..  2086
KBLV       .....V.K.SV........N...........N...............R..  2086
LBV        .....T.RSKK......ASPSPI..K.....S....A....M........  2086
LLEBV      ..R..TYR.RT....N.TSES..Y...S...MF........M...V.RLN  2085
SHIBV      ..R..T.K.KKF.....A.PS.I..K.S...S....A....M........  2086
MOKV       ..G..T.R.RK......ANSSPI..K.....SI.................  2086
VSV_NJ     F-R-VLKNEKWLQK.DCKGD-A.P.DSRLGDS.ANIGN...AWEL.RDGN  2047
```

```
Consensus  RLXGSXXDLSXEVEXHLXGYNRWIXXXDIRSR-------SSLLDYSCL--
RABV-Tha   .FI..IE..PG..AR..Q......TLE.....-------.........--  2128
ARAV       .FV..AK...K...R..R......NFN.....-------.........--  2127
ABLV       ..T..PV...K...K..R......TLN.VK..-------.........--  2128
BBLV       .FV.NTNE..K...K..R......NLN.....-------.......G.--  2127
WCBV       .VGE.GDII.RA.KSC.K......LMR.....-------T...E.....--  2127
DUVV       ..T.GPK...R.M.R..KS.....NFD.L...-------.......G.--  2127
TWBLV      ..A.NAK...R...KQ.KN.....SFE.L.I.-------T...E..A.--  2127
GBLV       ..A..HVE..R.I.K..K......TFN.....-------.........--  2127
EBLV1      ..V..AK...Q...K..KS.....NFS.L...-------.........--  2127
EBLV2      ..A..SN...K...K..K......SFD.....-------.........--  2127
IKOV       .INERGSKILD.MTTP.RT....LKIQ...H.-------T...E.WSN--  2126
IRKV       ..I.NAH...K...R..RS.....NFN.L...-------.........--  2127
KHUV       .FV..SS...E...K..R......KFH.....-------.........--  2127
KBLV       .FA..NN..AR...R..K......SLN.....-------....E.....--  2127
LBV        ..SCRPK..IR.T.AC.KN.....NMR.....-------T..M..C..--  2127
LLEBV      ..NEPKTR.MD..LVP.RS.....KLK.LKY.-------T...EFWSC--  2126
SHIBV      ..NSNPRE.LK...IY.K......TMR.V...-------T.....C..--  2127
MOKV       ..NC.PR.MLR.T.AC.RT..K..SIR.....-------T.IM..C..--  2127
VSV_NJ     KSEPFDSMVAETLTKSVDKSLS.RKISKSTGIPRLLNSDVD.V.Q.I.NV  2097
```

```
Consensus  ------------
RABV-Tha   ------------  2128
ARAV       ------------  2127
ABLV       ------------  2128
BBLV       ------------  2127
WCBV       ------------  2127
DUVV       ------------  2127
TWBLV      ------------  2127
GBLV       ------------  2127
EBLV1      ------------  2127
EBLV2      ------------  2127
IKOV       ------------  2126
IRKV       ------------  2127
KHUV       ------------  2127
KBLV       ------------  2127
LBV        ------------  2127
LLEBV      ------------  2126
SHIBV      ------------  2127
MOKV       ------------  2127
VSV_NJ     QIDIVENQAWQN  2109
```

**Figure S1. Comparison of amino acid sequences of the Large protein among the different species of lyssaviruses and VSV**. Alignment was performed with Clustal Omega in SnapGene version 8.0.3 on 19 different isolates. *Lyssavirus* Aravan (ARAV), EF614259 ; *Lyssavirus australis* (ABLV), AF081020; *Lyssavirus bokeloh* (BBLV), JF311903 ; *Lyssavirus caucasicus* (WCBV), EF614258 ; *Lyssavirus Duvenhage* (DUVV), EU293119 ; *Lyssavirus formosa* (TWBLV), MF472710 ; *Lyssavirus gannoruwa* (GBLV), KU244266 ; *Lyssavirus hamburg* (EBLV1)_MF187859. ; *Lyssavirus helsinki* (EBLV2), EF157977; *Lyssavirus ikoma* (IKOV), JX193798 ; *Lyssavirus irkut* (IRKV), EF614260; *Lyssavirus khujand* (KHUV), EF614261 ; *Lyssavirus kotalahti* (KBLV) LR994545 ; *Lyssavirus lagos* (LBV), EU293108; *Lyssavirus lleida* (LLEBV) , KY006983; *Lyssavirus mokola* (MOKV), EU293118 ; *Lyssavirus rabies (RABV),* EU293121 *; Lyssavirus shimoni (*SHIBV) , GU170201; *Vesiculovirus newjersey* (VSV NJ), JX121109.

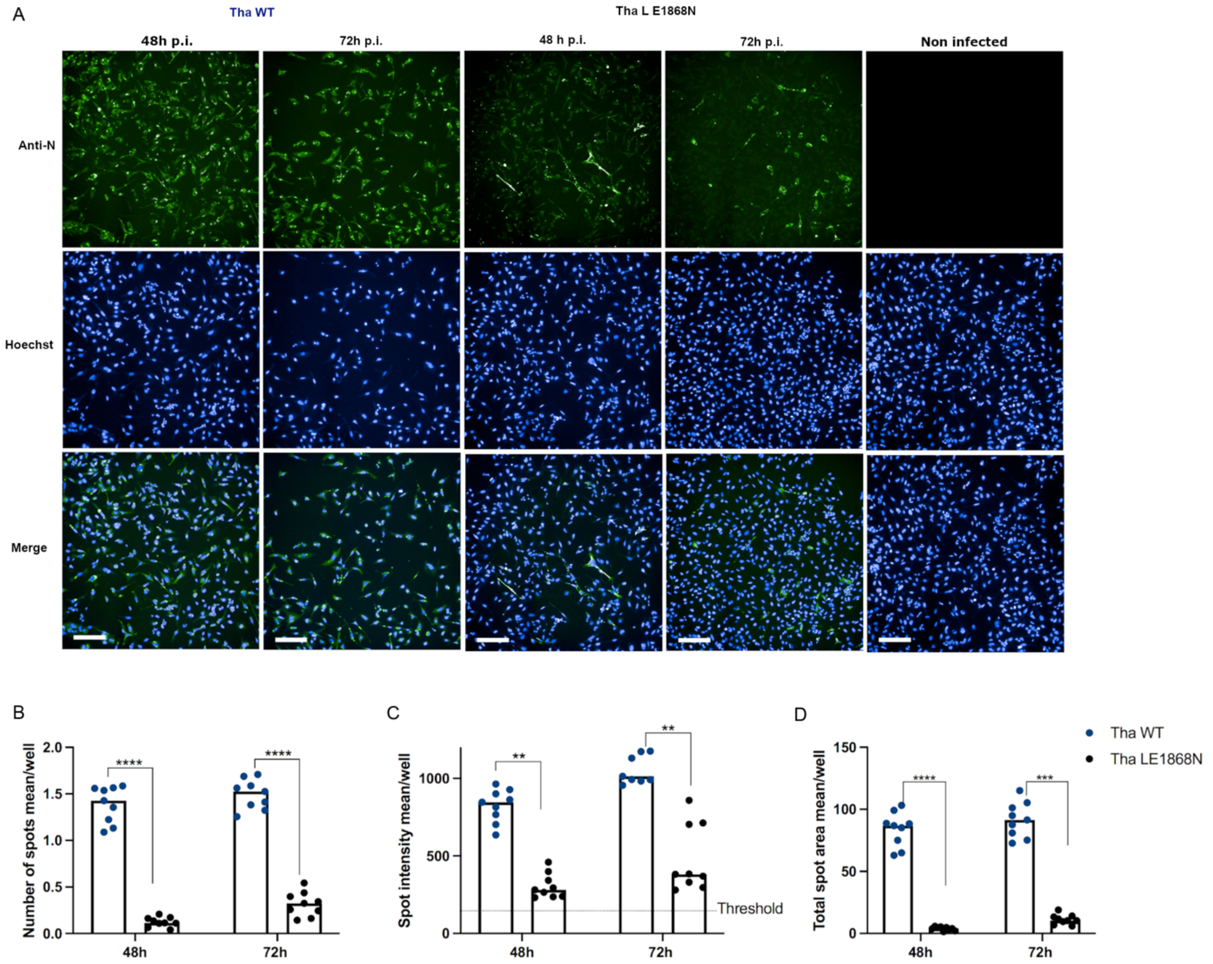


**Figure S2. A mutation on the residue E1869N in the MT domain of the L protein of RABV affects viral infection. (A)** Representative immunofluorescence pictures of SK-N-SH, upon infection with Tha and Tha-$L_{E1868N}$, respectively, at 48 h and 72h post-infection. Image acquisitions were performed on the automated confocal microscope Opera Phenix (Perkin Elmer) using the 10x objective. The nucleoprotein is marked in green, and the nuclei of the cells are in blue. **(B-D)** Characterisation of the spread of the mutated viruses. Image acquisitions were performed on the automated confocal microscope Opera Phenix (Perkin Elmer) using the 10× objective. Quantification of nucleoprotein expression (N) of SK-N-SH cells infected by Tha and Tha-$L_{E1868N}$ and Tha-$L_{T1681S}$ was performed at 48- and 72-hours post-infection. All experiments were performed three times (n=3) independently. Each dot represents imaging of one well of a 96-well plate (approx. 8 × $10^3$ cells/well). Three parameters were studied: the mean number of spots/well (B), the mean intensity of spots/well (C), and the mean total area of spots/well (D). Bars show mean ± SD. The percentages of $N^+$ cells were analysed using a mixed model with the replication factor as a random effect, followed by multiple comparisons corrected by Tukey's method (**** adjusted p-value < 0.000010, **

adjusted p-value < 0.00010, * adjusted p-value < 0.0010). If no p-value is indicated, no significant difference was observed.

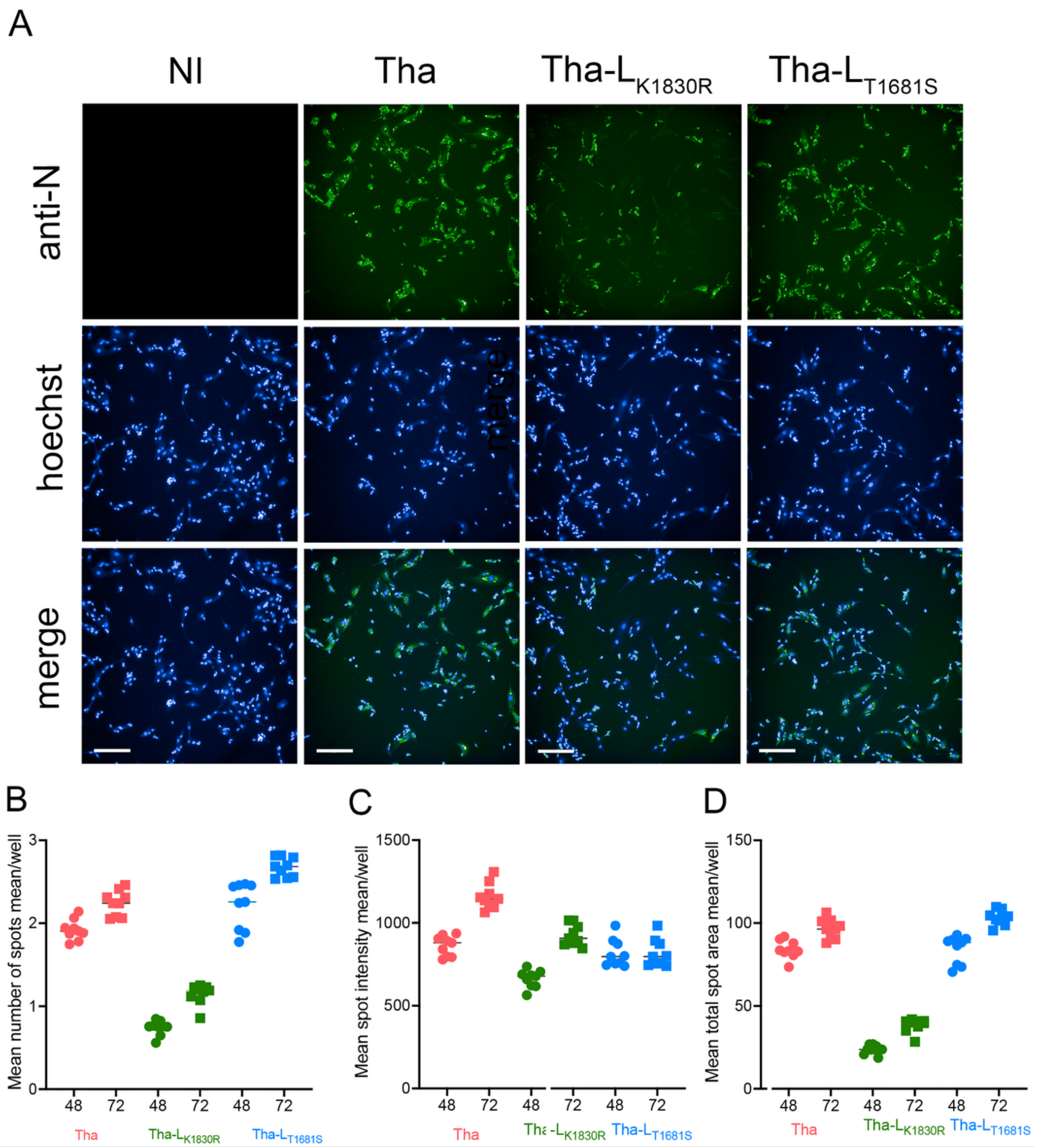


**Figure S3. A Mutation on the residue K1830R in the MT domain of the L protein of RABV affects viral infection compared to a mutation on T1681S. (A)** Representative immunofluorescence merge picture of SK-N-SH, upon infection with Tha, Tha-L$_{K1830R,}$ and Tha-L$_{T1681S,}$ respectively, at 48 h post-infection. Image acquisitions were performed on the automated confocal microscope Opera Phenix (Perkin Elmer) using the 10x objective. The nucleoprotein is marked in green, and the nuclei of the cells are in blue. **(B-D)** Characterisation of the spread of Tha, Tha-L$_{K1830R,}$ and Tha-L$_{T1681S}$ on SK-N-SH cells. Image acquisitions were performed on the automated confocal microscope Opera Phenix (Perkin Elmer) using the 10x objective. Quantification of nucleoprotein expression (N) of SK-N-SH cells infected by Tha, Tha-L$_{K1830R,}$ and Tha-L$_{T1681S}$ was performed at 48- and 72-hours post-infection. Nucleoprotein quantification was achieved by measuring FITC-labelled anti-nucleoprotein immunofluorescence in the channel 480/500-550. All experiments were performed three times (n = 3) independently. Each dot represents imaging of one well of a 96-well plate (approx. 8 × $10^3$ cells/well). Three parameters were studied: the mean number of spots/well (A), the mean intensity of spots/well (B), and the mean total area of spots/well (C). Bars show mean ± SD. The percentages of $N^+$ cells were analysed using a mixed model with the replication factor as a random effect, followed by multiple comparisons corrected by Tukey's method (**** adjusted p-value < 0.000010, ** adjusted p-value < 0.0001, * adjusted p-value < 0.001). If no p-value is indicated, no significant difference was observed.

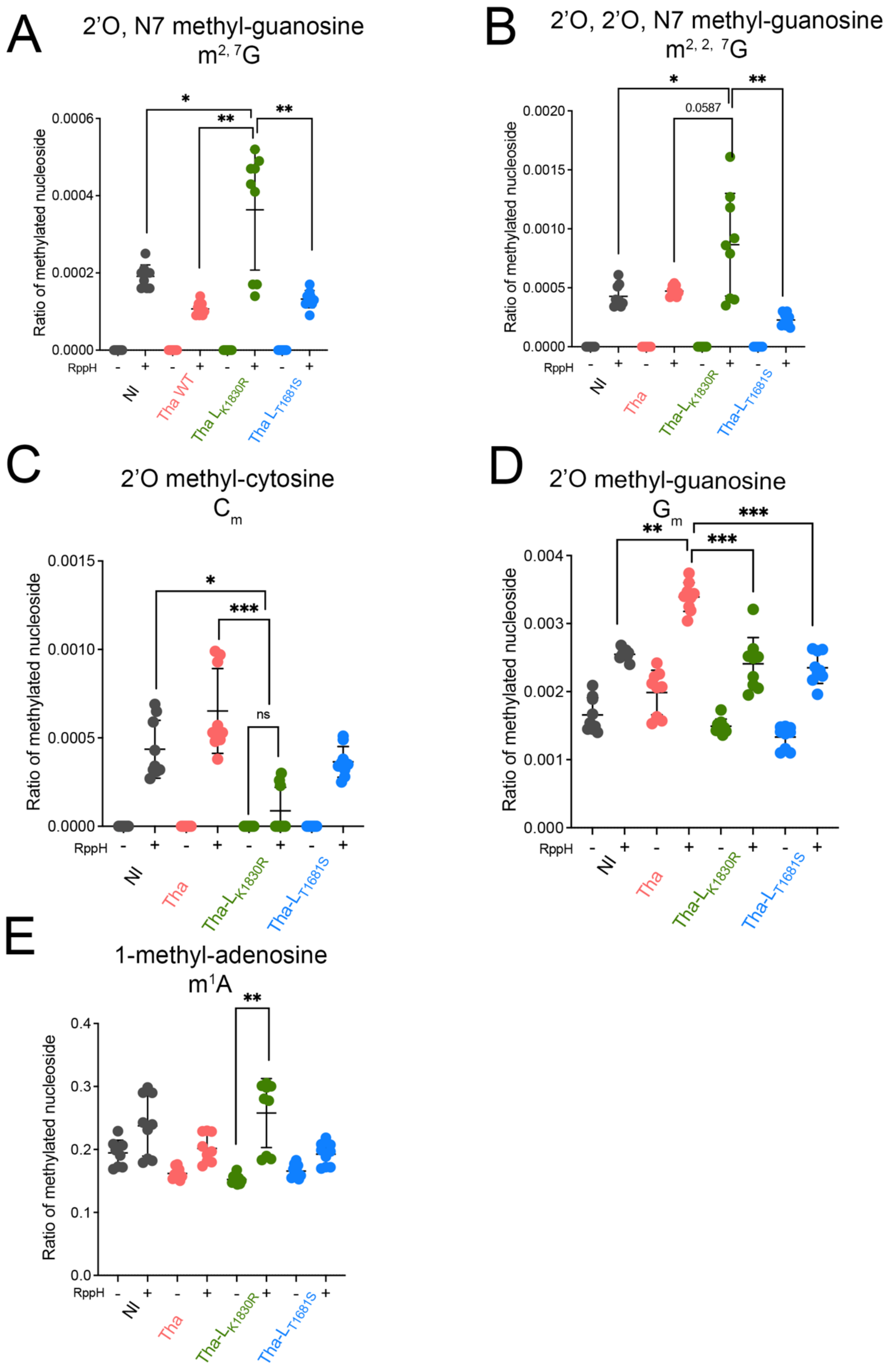


**Figure S4. Quantitative LC-MS/MS measurements of 5′ terminal and internal modifications in poly(A) mRNA** extracted from SK-N-SH cells infected with Tha, Tha-$L_{K1830R}$, and Tha-$L_{T1681S}$. **(A-B)** Levels of m27G and m227G modifications on the cap structure of purified mRNA from infected SK-N-SH cells (*n* = 3). The levels of nucleoside modifications were normalized to the total molar amount of canonical nucleosides N (N = C + U + G + A) **(C-D).** Level of 2'O-methylation on C or G nucleoside, in the internal position of mRNA from infected SK-N-SH cells (*n* = 3). The levels of nucleoside modifications were normalized to the total molar amount of canonical nucleosides N (N = C + U + G + A). E) Level of N1-methylation on A nucleoside, in the internal position of mRNA from infected SK-N-SH cells (*n* = 3). Results are average over the levels of repbio. P-value adjustment was calculated by the Tukey method for comparing a family of 4 estimates. Significant results are indicated by star: *: $p<0.05$; **: $p<0.01$; ***: $p<0.001$; ****: $p<0.0001$.

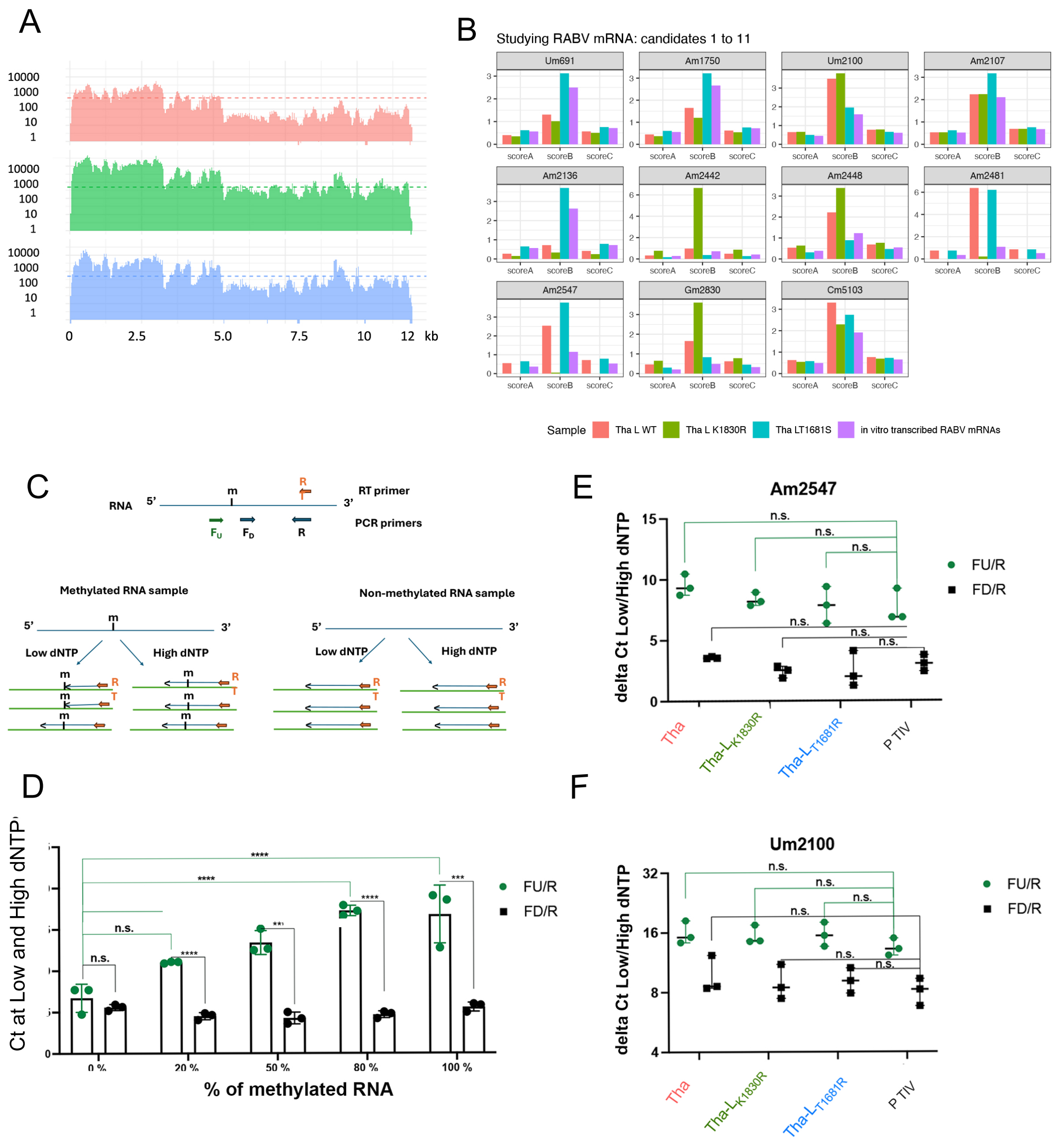


**Figure S5: Detection of 2'O methylation in the internal position of mRNA from infected SK-N-SH cells using RibometSeq. (A)** Comparison of coverage obtained for the sequencing of 3 different infections. **(B)** RiboMethScores obtained for the 11 methylated candidates. 5’ and 3’-ends coverage were normalized with DESeq2, ribomethylated positions coverages and scores are plotted using ggplot2 under R version 4.0.3 using ggplot2. **(C)** General principle of the RTLQ technique used to validate the 2’O-methylated candidates. Reverse transcription was performed at low and high dNTP concentrations. If the RT encounters methylation, the cDNA will stop at the methylation site. Two PCR systems ($F_U$/R and $F_D$/R) were used to amplify both cDNA, which contain a potential site of methylation. The Ct value obtained by SYBR Green amplification was recorded for the low and the high dNTP concentration. **(D)** Calibration of the RTLQP technique on the Um2100 system. Mixtures of synthetic RNAs methylated or not on the Um2100 residue were mixed according to different ratios (0, 20, 50, 80, and 100% of methylated RNA). After cDNA was synthesized with low and high dNTP concentrations, QPCR was performed with both primer systems. Ct value obtained for each mixture on both systems is plotted: in green, the $F_U$/R system, and in black, the $F_D$/R system. **(E-F)** Validation of the positions of Am2547 and Um2100 identified by RiboMethSeq. The previously extracted mRNA from infected cells was used as a matrix for cDNA synthesis with both low and high dNTP concentrations. Ct values obtained for each condition are indicated on the graphs. Statistical differences obtained between the TIV (without methylation) and the mRNA from infected cells were calculated using multiple comparisons corrected by Tukey’s method. ns: non-significant results, *: $p<0.05$; **: $p<0.01$; ***: $p<0.001$; ****: $p<0.0001$.

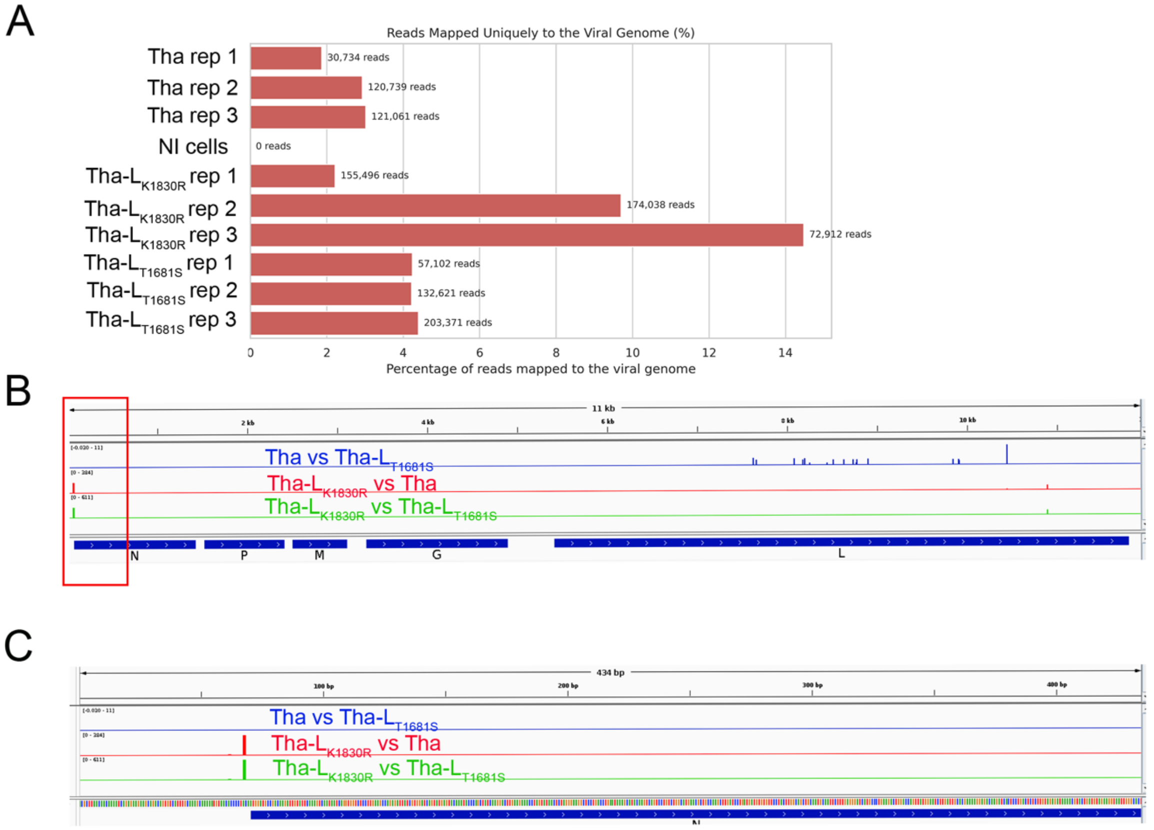


**Figure S6: Detection of m6A methylation in the internal position of mRNA from infected SK-N-SH cells using Direct-RNA sequencing. (A)** Comparison of coverage obtained for the sequencing of the different infections. All conditions were sequenced in triplicate. The percentage of reads mapped to the viral genome is indicated on the x-axis. **(B)** Distribution of differentially methylated positions between each comparison pair. Methylation rates were computed using modkit pileup for each replicate, and differential modification analysis was performed using modkit dmr pair for the three pairwise comparisons between samples (Tha-L$_{K1830R}$ vs Tha in blue, Tha vs Tha-L$_{T1681S}$ in green, and Tha-L$_{K1830R}$ vs Tha-L$_{T1681S}$ in red). Green and red bars correspond to position 68 on the Tha genome, which is significantly more methylated in Tha-L$_{K1830R}$ conditions than the two other infections **($p$-value $< 0.05$**, with a high confidence score, and an **effect size > 30%). (C) It** is a zoom of the part framed in red on part B.

**Table S1**: Sequence of the primers used for mutagenesis and RT-Low dNTP PCR (RTLP) approach. Sequences are in 5'->3' orientation.

| | | |
|---|---|---|
| **Primers for mutagenesis** | | |
| **K1686R** | K1686R-for | AACTGGCGCCCATTATAGACTCAAGCCCATTCTAG |
| | K1686R-rev | TGCCCACTGGACAACTCTCAAGCCCGAGATAAG |
| **D1798 N** | D1798N-for | TGACCTAATCATTTGTAACGCAGAAGTCACTGACA |
| | D1798N-rev | TAAGACATGTTCACCTGTTTTTGG |
| **K1830R** | K1830R-for | TCTCTATCTGGTCTTCCGAACTTACGGAACCATGC |
| | K1830R-rev | GGACCATCTATGGATAATGCAAAGTCTGAC |
| **E1868N** | K1686R-for | CTCGTCTTTTTCATCCAACCTGTACCTCAGATTCTC |
| | K1686R-rev | GTCATTTGGGTTATAAATCCTGTGACTGAAGG |
| **T1681S** | T1681S-for | AGTTGTCCAGTGGGCATCCGGCGCCCATTATAAGC |
| | T1681S-rev | CTCAAGCCCGAGATAAGAGGGTTTTGG |
| **Primers for RT-Low dNTP PCR (RTLP)** | | |
| **Um2100** | Um2100 For | TGG TCCTCCAGCTCTCGA AT |
| | Um2100 FD | CTGTAGAGGCAGAAATAGCC |
| | Um2100 rev | TTTCTTGGAAAAGCTCTCAGC |
| **Am2547** | Am2547 For | TTTTTTTCATGTCAGGTCCG |
| | Am2547 FD | AGCCCTCCCCTGTATCGG |
| | Am2547 rev | CAGGAGGAGGCAGCCACAGGTC |
| **Am1909** | RT Am1909 | GCCTCCATTCGGGCTTTCAG |
| | Fu Am1909 | CTGGTCACAAACCGTGGAGG |
| | FD Am1909 | CGGGAAGGTCTTTGGAAGAT |
| **Primers for *in vitro* transcription** | | |
| **P2RZ-N** | T7-PromN 59For | ACGACTCACTATAGGAACACCCCTACAATGGATG |
| | N1483HDV Reve | ACCATGGCTAGCGTTTTTTTCATGGGAGATGTACACT |
| **P2RZ-P** | T7-PromP1486 For | ACGACTCACTATAGGAACACCCCTCCTTTTGAACCATCCC |
| | P2475-HDV Rev | ACCATGGCTAGCGTTTTTTTCATGTCAGGTCCGGAAC |
| **P2RZ-M** | T7-PromM2481For | ACGACTCACT ATAGGAACACCACTGATAAAATGAA |
| | M3283HDV Rev | ACCATGGCTAGCGTTTTTTTCACATCCAAGAGGCT |
| **P2RZ-G** | T7- G3301 For | ACGACTCACTATAGGAAGACTCAAGGAAAGATGAT |
| | G 5347-HDV Rev | ACCATGGCTAGCGTTTTTTTCTCGACTGAAAAGCG |
| **P2RZ-L** | T7-Prom-L 5370 For | ACGACTCACTATAGGAACACTTCTCATCTTCAGGC |
| | L 11848-HDV Rev | GGACCATGGCTAGCGTTTTTTTCACGGTACACTGT |

**Table S2:** Comparison of coverage obtained for the sequencing of the different infections using RibomethSeq.

| Sample name | Total reads | Reads mapped on Tha sequence | Rate on Tha sequence (%) |
|---|---|---|---|
| Tha-R1 | 24805216 | 536571 | 2.16 |
| Tha-R2 | 14714326 | 227073 | 1.54 |
| Tha-R3 | 22929003 | 486184 | 2.12 |
| Tha-$L_{K1830R}$-R1 | 16800673 | 3718869 | 22.14 |
| Tha-$L_{K1830R}$-R2 | 26542939 | 5484522 | 20.66 |
| Tha-$L_{K1830R}$-R3 | 30645333 | 6312837 | 20.6 |
| Tha-$L_{T1681S}$-R1 | 48277820 | 1475137 | 3.06 |
| Tha-$L_{T1681S}$-R2 | 25356951 | 964928 | 3.81 |
| Tha-$L_{T1681S}$-R3 | 31733252 | 1011171 | 3.19 |
| Mix IVT1 | 27931419 | 21136193 | 75.11 |
| Mix IVT2 | 12459532 | 9358359 | 75.11 |
| Mix IVT3 | 18826509 | 14227189 | 75.57 |
